# HANSEN: An Integrated Structural and Functional Proteome Resource for Structure-Guided Drug Discovery in *Mycobacterium leprae*

**DOI:** 10.64898/2026.08.03.741541

**Authors:** Sundeep Chaitanya Vedithi, Rebecca Rees, Sony Malhotra, Asma Munir, Modestas Matusevicius, Ali F. Alsulami, Christopher A. Beaudoin, Kiran Sai Sunkara, Madhusmita Das, Tom L. Blundell, Rodrigo Andres Floto

## Abstract

Leprosy remains a leading infectious cause of preventable disability, yet its causative agent, *Mycobacterium leprae (M. leprae)*, is structurally under-characterised. Only ten Protein Data Bank (PDB) entries represent seven of its 1,603 protein-coding genes. We present HANSEN, a proteome-wide structural and functional resource for *M. leprae*. Monomeric and oligomeric models were generated with AlphaFold 3, Boltz-1, Boltz-2 and Chai-1, and annotated with per-residue confidence, predicted aligned error and, for assemblies, interface confidence. Ligand-binding pockets were predicted with AF2BIND, P2Rank and fpocket, template-derived ligands were modelled within oligomeric complexes, residue-level B-cell epitope propensity was estimated with DiscoTope-3.0, and gene essentiality was transferred from *Mycobacterium tuberculosis* transposon-sequencing labels. These features are integrated in a relational database with interactive visualisation and combined into a calibrated Target Priority Score that ranks all 1,603 proteins into four tiers and recovers established antimycobacterial targets. HANSEN (https://hansen-leprosy.medschl.cam.ac.uk/home) provides a practical basis for target prioritisation and structure-guided drug discovery in leprosy.

## 1. INTRODUCTION

Leprosy, or Hansen’s disease, remains an important mycobacterial infection of public-health concern, particularly in tropical and subtropical regions. A total of 172,717 new cases were reported worldwide in 2024 (*1*). The disease is caused primarily by an acid-fast bacillus *Mycobacterium leprae* (*M. leprae*) and, in some cases, by *Mycobacterium lepromatosis*. These organisms have a pronounced tropism for Schwann cells of the peripheral nerves and for dermal cell populations. Untreated or inadequately treated infection can lead to chronic neuropathy, progressive demyelination, impaired sensory and motor function, cutaneous lesions, tissue damage and enduring social stigma (*2*). Standard treatment is multidrug therapy (MDT), which combines dapsone, rifampicin and clofazimine for six to twelve months according to whether disease is paucibacillary or multibacillary (*3*). Although MDT has substantially reduced leprosy prevalence, the prolonged regimen burdens patients and may reduce adherence. It does not reverse established nerve damage, and axonal loss may continue despite antimicrobial treatment (*4*). Resistance to one or more MDT components also threatens the durability of current regimens (*5*). Therapeutic development for *M. leprae* has progressed much more slowly than for *M. tuberculosis*, and many leprosy candidates have been repurposed from the tuberculosis pipeline (*6*). The persistent disease burden, treatment limitations, inadequate protection against nerve damage and emergence of resistance therefore create an urgent need for new pharmacological strategies and molecular targets.

A major obstacle to *M. leprae* research and drug development is the organism’s experimental intractability. *M. leprae* is an obligate intracellular pathogen that cannot be propagated in axenic culture, removing a fundamental tool for antimicrobial discovery. Research instead relies on *in vivo or ex vivo* systems, including mouse footpad (*7*) and armadillo infection models (*8*). These systems are expensive, labour-intensive, slow and unsuitable for high-throughput studies. Consequently, the structural proteome of *M. leprae* remains poorly understood. Of the 1,603 protein-coding open reading frames, only 10 PDB entries represent seven unique proteins, compared with approximately 1,800 experimentally determined structures for *M. tuberculosis* (*9*). Many *M. leprae* orthologues of pharmacological or mechanistic importance, including enzymes involved in folate metabolism, RNA polymerase and DNA gyrase, lack experimentally determined three-dimensional structures. This gap restricts rational drug design, ligand docking, mechanistic interpretation and target-based functional investigation.

Recent advances in computational structural biology provide a practical way to overcome many of these barriers. Modern artificial intelligence (AI) based prediction methods can generate useful three-dimensional models where experimental structures are unavailable. The AlphaFold Protein Structure Database (AFDB) now provides predicted models for 1,602 sequences in the *M. leprae* TN reference proteome (*10*). These models substantially expand structural coverage but are limited to monomeric proteins and therefore do not capture higher-order organisation in the form of homo- and hetero-oligomeric assemblies, restricting insights into small molecule binding and protein function. In tuberculosis, CHOPIN demonstrated the value of integrating proteome-scale structural models with functional information in a dedicated web resource (*11*).

Computational structural modelling can bridge the gap between sequence and structural information in *M. leprae*. Accessible web resources also broaden the use of structural information among researchers without specialist modelling expertise or extensive computational infrastructure. Comparative modelling of the *M. leprae* RNA polymerase complex showed how clinically observed *rpoB* mutations could affect rifampicin binding, protein stability and interactions with the growing RNA transcript (*12*). Computational saturation mutagenesis then evaluated 21,394 possible substitutions in the RNA polymerase β-subunit and identified changes predicted to affect stability and inter-molecular interactions (*13*). HARP database extended this approach across RNA polymerase, dihydropteroate synthase and DNA gyrase, providing predicted structural effects for possible 80,902 missense substitutions (*14*). These studies illustrate how computational structures can generate mechanistic hypotheses and prioritise mutations for experimental investigation.

Here, we present HANSEN, a dedicated web resource providing predicted three-dimensional structures and associated annotations for almost the entire *M. leprae* proteome. To our knowledge, it is the first proteome-wide *M. leprae*-specific platform to integrate proteome-wide structural models with functional features. To reduce reliance on one modelling approach and permit cross-method comparison, HANSEN combines AlphaFold 3(*15*), Boltz-1(*16*) and Chai-1(*17*) models generated through the version of ABCFold(*18*) available at the time with a separately generated Boltz-2 model set (*19*). Boltz-2 builds on Boltz-1 through expanded structural training data, including experimental and molecular-dynamics ensembles, improved representation of conformational variability and modest gains in structural accuracy across several biomolecular modalities. It additionally introduces joint prediction of protein–ligand structure and binding affinity. Template information was also used to model putative protein assemblies and derive oligomeric states. Each model is evaluated with residue-level and inter-residue confidence measures. The annotations cover protein domains, catalytic features, predicted ligand-binding pockets, information related to ligand and cofactors in oligomers, B-cell epitope propensity, pathways, gene-essentiality derived from homologues in *M. tuberculosis* and an interface to process sequencing data to detect antimicrobial resistance (AMR). This integrated framework is intended to support target prioritisation, virtual screening and subsequent experimental investigation.

## 2. RESULTS

### 2.1 A proteome-wide structural atlas for *M. leprae*

HANSEN provides broad structural coverage across the *M. leprae* proteome, addressing the scarcity of experimentally determined structures. The dataset contains 1,603 protein records, of which 1,599 carry a UniProt accession and four are modelled from genomic translations that have no UniProt entry (Supplementary Note SN1). Of these, 669 are reviewed UniProt entries and 1,205 have Gene Ontology annotation (*20–22*), whereas only seven have experimental three-dimensional structures (Table 1). By integrating AI based and template-informed modelling, HANSEN provides predicted structures for most proteins. Users can inspect modelling method, confidence measures, PAE, predicted pockets, ligand features and functional annotations. The Query page enables targeted retrieval when a specific protein is known. Users can search using a UniProt accession, *M. leprae* gene name, a protein name, a ligand identifier such as a code, name, SMILES or PubChem CID, or a full or partial amino acid sequence. Identifier and name-based searches support exact, prefix and partial matching with autocomplete. Sequence queries are normalised and analysed using local alignment. A successful search opens the corresponding protein page, where users can examine structural models, predicted pockets, ligands and associated annotations. In contrast, the Gene Ontology (GO)Browser supports exploratory navigation by displaying GO terms as an interactive sunburst across the biological process, molecular function and cellular component namespaces. This allows users to identify groups of functionally related proteins based on ontology terms rather than specific identifiers. Selecting a term returns the associated *M. leprae* proteins in a table, with each result linked to its protein page using UniProt accession, thereby enabling database-wide functional categories to be explored at the level of individual protein structures, annotations and supporting functional information.

**Table 1.** HANSEN database composition:

| Resource metric | Value |
| --- | --- |
| Proteome records | 1,603 |
| Reviewed (UniProt) entries | 669 |
| Entries with Gene Ontology annotations | 1,205 |
| Entries with experimental 3D structures | 7 (10 structures in PDB for 7 unique proteins) |
| Homo-oligomeric structural model records | 528 (506 Boltz-2 + 22 AlphaFold3 homooligomeric assemblies) |
| Hetero-oligomeric structural model records | 198 (heterooligomeric assemblies) |
| Registered structural model records | 7,122 |
| Modelling methods integrated | AlphaFold 3, Boltz-1, Boltz-2, Chai-1 |
| Ranked protein models | 1,603 |
| High-priority protein targets (score $\geq 67$) | 50 |
| Strong candidate protein targets (score 61–67) | 125 |
| Protein models with predicted pockets | 1,603 |

### 2.2 Integration of Multiple Modelling Methods

HANSEN incorporates structural models generated by AlphaFold 3, Boltz-1, Boltz-2 and Chai-1. This multi-method approach allows users to compare predictions and assess method-specific confidence. The statistics dashboard summarises registered model records by method, while protein pages allow individual models to be loaded and inspected. The highest-confidence monomeric models illustrate the quality attainable across the atlas (Table S1).

### 2.3 Oligomeric Assembly Representation

HANSEN extends beyond monomeric structures by incorporating predicted oligomeric assemblies classified by chain count and assembly type. The current database contains 726 oligomeric model records for 720 proteins, comprising 528 homo-oligomeric and 198 hetero-oligomeric records. A protein may belong to more than one modelled assembly, so the number of records exceeds the number of proteins. The acyl carrier protein AcpM (ML1654), for example, is modelled both within the EmbA–EmbB arabinosyltransferase complex (template PDB 7BVF, three chains, co-folded) and in a two-chain assembly with the transporter MmpL4 (template PDB 9U51, built by superposition). Template PDBs for assemblies are listed in Table S2. For oligomeric assemblies, the predicted aligned error (PAE) matrix shows how confidently the chains are positioned relative to one another. Low off-diagonal values support the placement of one chain against another, whereas high values indicate uncertainty in their relative orientation.

Representative homomers include inorganic pyrophosphatase Ppa (ML0210), modelled as a homohexamer, and dihydroneopterin aldolase FolB (ML0225) (Fig. 2a), modelled as a homooctamer. Heteromeric examples include the ML0596–ML0597 cysteine-desulfurase complex and the ML0112–ML0114 ABC-transporter complex (Fig. 2b), both modelled as hetero-tetramers (Table S2).

**Fig. 1.**
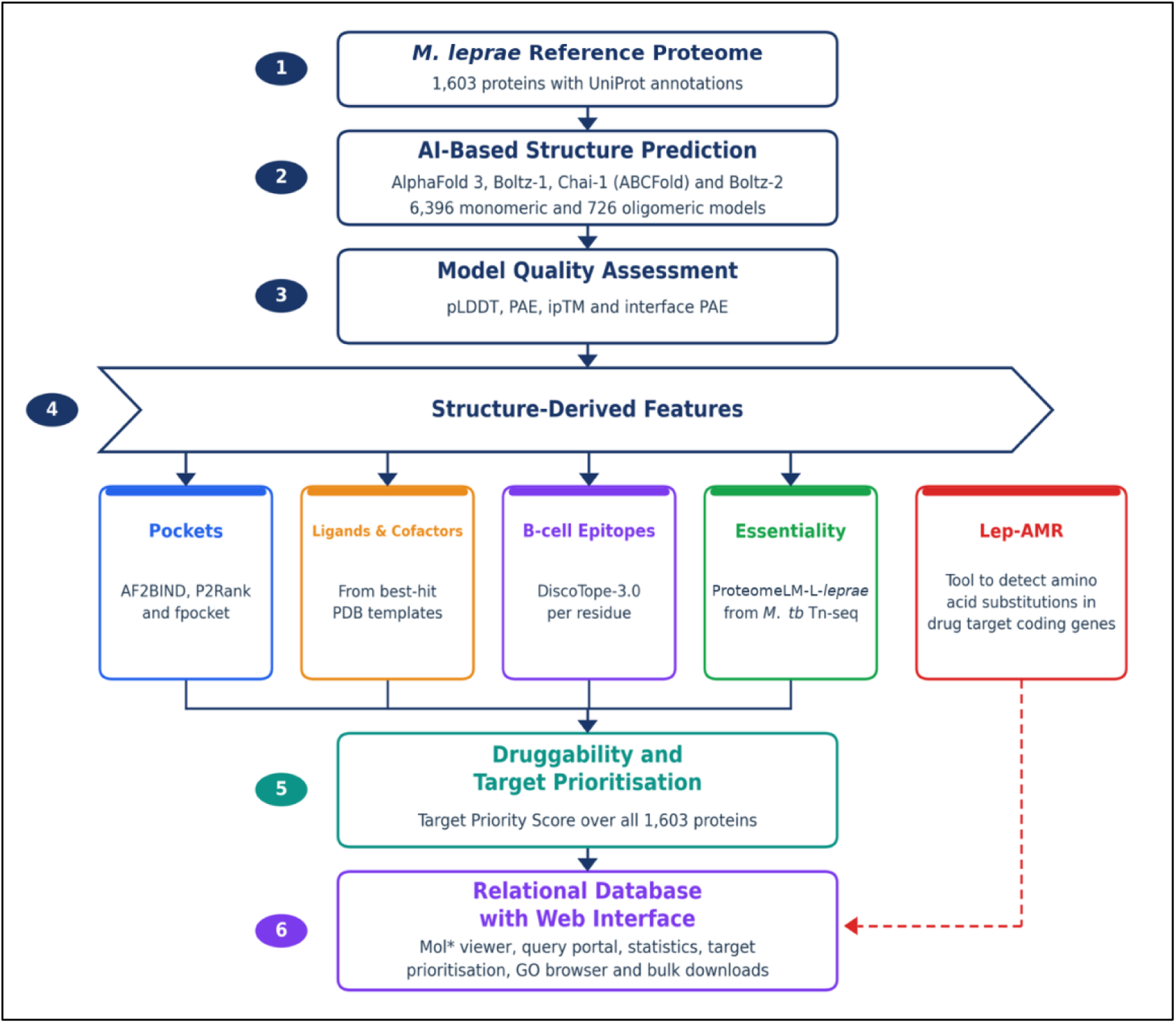
Overview of the HANSEN workflow. The *M. leprae* protein set (1) is modelled with complementary AI based methods through ABCFold and a separate Boltz-2 workflow (2). Monomeric models are generated by multiple structure predictors (AlphaFold3, Boltz, Boltz2 and Chai). Oligomeric assemblies both homomeric and heteromeric are produced predominantly by Boltz2 co-folding (n=564 of 726oligomers). Where direct co-folding is unsuitable due to the size of the monomers, assemblies are instead reconstructed by superimposing the monomeric models onto experimental template assemblies (structure superimposition; n=140 oligomeric model records from 121 distinct models), and a set of homomers are also modelled with AlphaFold3 (n=193), of which the 22 without a Boltz-2 counterpart are retained as distinct assemblies. All oligomeric models are checked for chain geometry, inter-chain overlap, chain completeness and ligand placement. Model quality is assessed using predicted local distance difference test (pLDDT) scores and predicted aligned error (PAE) values (3). Structure-derived features which include predicted pockets from AF2BIND, P2Rank and fpocket, template-inherited ligands and cofactors, B-cell epitope propensity and ProteomeLM-derived essentiality are computed on the models (4). Druggability and the Target Priority Score (5) are calculated from those structural features and Lep-AMR is shown as a separate module for detecting substitution mutations in drug target coding genes. Users can access the resource through a web interface with a Mol* viewer for molecular visualisation, query portal, statistics dashboard, target-prioritisation browser and bulk-download functions for tables (6).

**Fig. 2.**
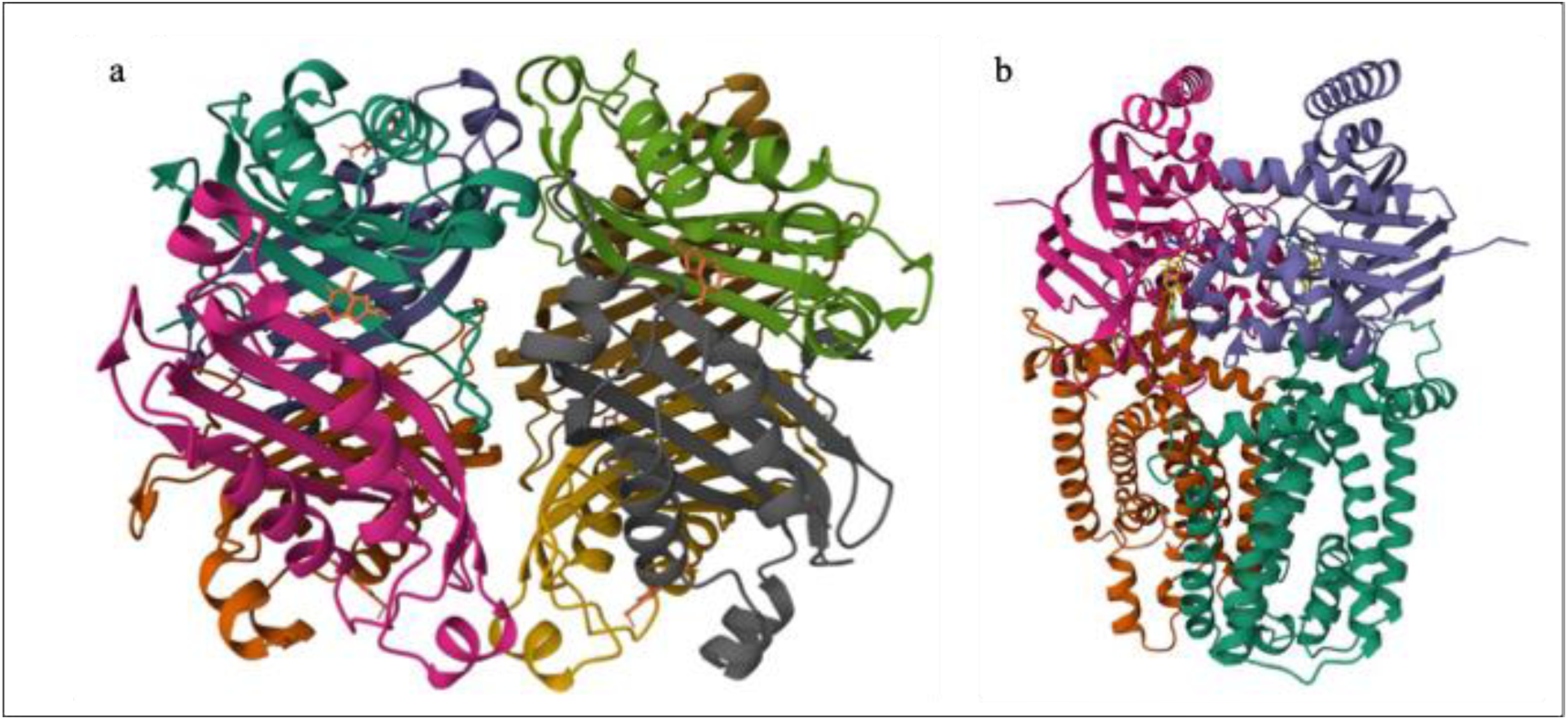
(a) Homo-octameric structure of dihydroneopterin aldolase FolB (ML0225) bound to B55. (b) Heterotetrameric structure of ML0112–ML0114 ABC-transporter complex bound to ATP.

Boltz-2 ranking scores were available for 564 of the 726 oligomeric model records in Table S2. Of the remaining 162 records, 140 were built by rigid-body superposition of monomeric models onto template assemblies and were not evaluated with Boltz-2. The remaining 22 records were generated with AlphaFold3 and therefore had AlphaFold3 ranking scores instead. In total, 586 records had a ranking score: 564 from Boltz-2 and 22 from AlphaFold3. Among the Boltz-2 models, the mean score was 0.821 for 480 homomers (median 0.836; range 0.459– 0.972) and 0.814 for 84 heteromers (median 0.838; range 0.604–0.956). The remaining 22 homooligomers modelled by AlphaFold3 has a mean ranking score of 0.711 (median: 0.762; range: 0.234–0.970).

As the ranking score does not measure interface confidence, ipTM and interface predicted aligned error were calculated for each oligomeric record and reported in Table S2. Interface confidence was available for 559 of the 726 records. Of the remainder, 140 were built by rigid-body superposition of monomeric models onto template assemblies and 22 were generated with AlphaFold 3, so no Boltz-2 co-folding confidence exists for them, and for the remaining 5 records the co-folding confidence output was not retained. The mean ipTM was 0.719 and the median was 0.813. Of these records, 283 had an ipTM of at least 0.8, while 124 had an ipTM below 0.5.

Heteromeric assemblies had a slightly higher mean ipTM than homomeric assemblies, at 0.758 compared with 0.712. Only 8 of the 84 heteromers had an ipTM below 0.5, compared with 116 of the 475 homomers. The mean interface predicted aligned error was 8.7 Å for the same 559 records with 9.0 Å for the homomers and 7.3 Å for the heteromers. Assemblies with an ipTM below 0.5 are considered to have unsupported interfaces and are flagged accordingly in the resource.

### 2.4 Binding Site Predictions

HANSEN provides binding-site predictions for the 1,603 proteins of the *M. leprae* proteome through three methods. The fpocket(*23*) detects cavities from protein geometry by clustering alpha-spheres and reports 6,950 cavities across all 1,603 proteins. P2Rank(*24*) scores surface pockets for ligandability with a machine-learning model and reports 40,500 pockets across 1,544 proteins; P2Rank was applied to every model, but 59 predominantly small proteins (median length 87 residues, against 291 for the remainder) have no ligandable cluster and were dropped during P2Rank filtering; however, these are covered by fpocket. A consensus procedure integrates the two, assigning each pocket a consensus score, a volume and a druggability score, and reports 47,194 pockets across all 1,603 proteins. Because pockets are predicted independently on each structure model of a protein (fpocket on 6,920 models, P2Rank on 6,039), these totals are aggregated over models and do not correspond to distinct binding sites. AF2Bind(*25*) scores each residue for its probability of contacting a small molecule from AlphaFold representations and reports 21,204 candidate binding-site residues across all 1,603 proteins, of which 17,223 exceed a probability of 0.5. The full per-residue score distribution is provided in the Table S4. These outputs define the residues and surface regions used as the search space in molecular docking and virtual screening, and provide pocket volume, druggability and per-residue binding scores for ranking targets. Of the 1,603 proteins, 7 have an experimentally determined structure, for which predicted pockets are also computed (Table S4).

### 2.5 Interactive Structural Exploration

The Mol* viewer(*26*) provides an interactive environment for inspecting HANSEN structures (Figs. S7a and S7b). Users can load predicted monomeric and oligomeric models, colour them by pLDDT, focus on ligands, inspect pockets, highlight residues and load PAE heatmaps on demand. The structure loading strategy improves responsiveness for large oligomers, and feature tables are linked to the viewer so that residues, pockets and confidence parameters can be explored interactively. Screenshots of the protein page and the target-prioritisation browser are provided as Figs. S7 and S8.

### 2.6 B-cell Epitope Propensity

HANSEN includes structure-based B-cell epitope predictions from DiscoTope-3.0 (*27*), computed on the AlphaFold3 and Boltz-2 models of all 1,603 proteins, which score 1,078,718 residues and flag 216,853 of them (20.1%) as predicted epitope residues. Each residue carries an epitope propensity score and its relative solvent accessibility, and can be coloured onto the structure, allowing users to locate surface regions of elevated propensity and to assess whether these coincide with accessible loops, predicted pockets or other functionally relevant surfaces (Table S5).

The proportion of flagged residues is almost identical across the predicted localisation groups. Among the 1,600 proteins that carry a UniProt localisation annotation, it is 20.00% for the 1,245 proteins with neither a transmembrane segment nor a signal peptide, 20.47% for the 310 proteins with a transmembrane segment and 20.36% for the 45 proteins with a signal peptide. This consistency shows that the flagging rate reflects the default DiscoTope 3.0 threshold rather than differences in antigen biology. Although *M. leprae* is an obligate intracellular pathogen, bacillary lysis in the circulation may release intracellular proteins. Once released, these antigens can be recognised by B-cell receptors and circulating antibodies. Proteins without a transmembrane segment or signal peptide should therefore not be excluded solely based on predicted localisation. Their serodiagnostic value depends on antigen release, epitope integrity and recognition by antibodies in patients with leprosy.

### 2.7 Essentiality and Target Prioritisation

HANSEN integrates essentiality prediction and AI-assisted target prioritisation to support translational interpretation. Rather than assigning a rigid binary classification of essentiality, the platform combines essentiality with other features and acknowledges the effects of orthology, model assumptions and organism-specific biology (Table S7). The approach (described in Methods 4.10) ranks 1,603 proteins and identifies 50 high-priority, 125 strong, 513 moderate and 915 exploratory candidates (Fig. S8, Table S6). The ranking supports selection for *in-silico* screening, biochemical validation or medicinal-chemistry follow-up.

#### 2.7.1 Cross-species essentiality prediction with ProteomeLM-L-*leprae*

ProteomeLM-L-*leprae* model was applied to the *M. leprae* proteome, assigning each protein a calibrated essentiality probability and one of four classes. The classifier was trained here on ProteomeLM-L contextual embeddings (28). Of the 1,603 proteins, 90 were classified as essential, 79 as likely essential, 101 as uncertain and 1,333 as non-essential (Fig. 3, Table 2).

**Fig. 3.**
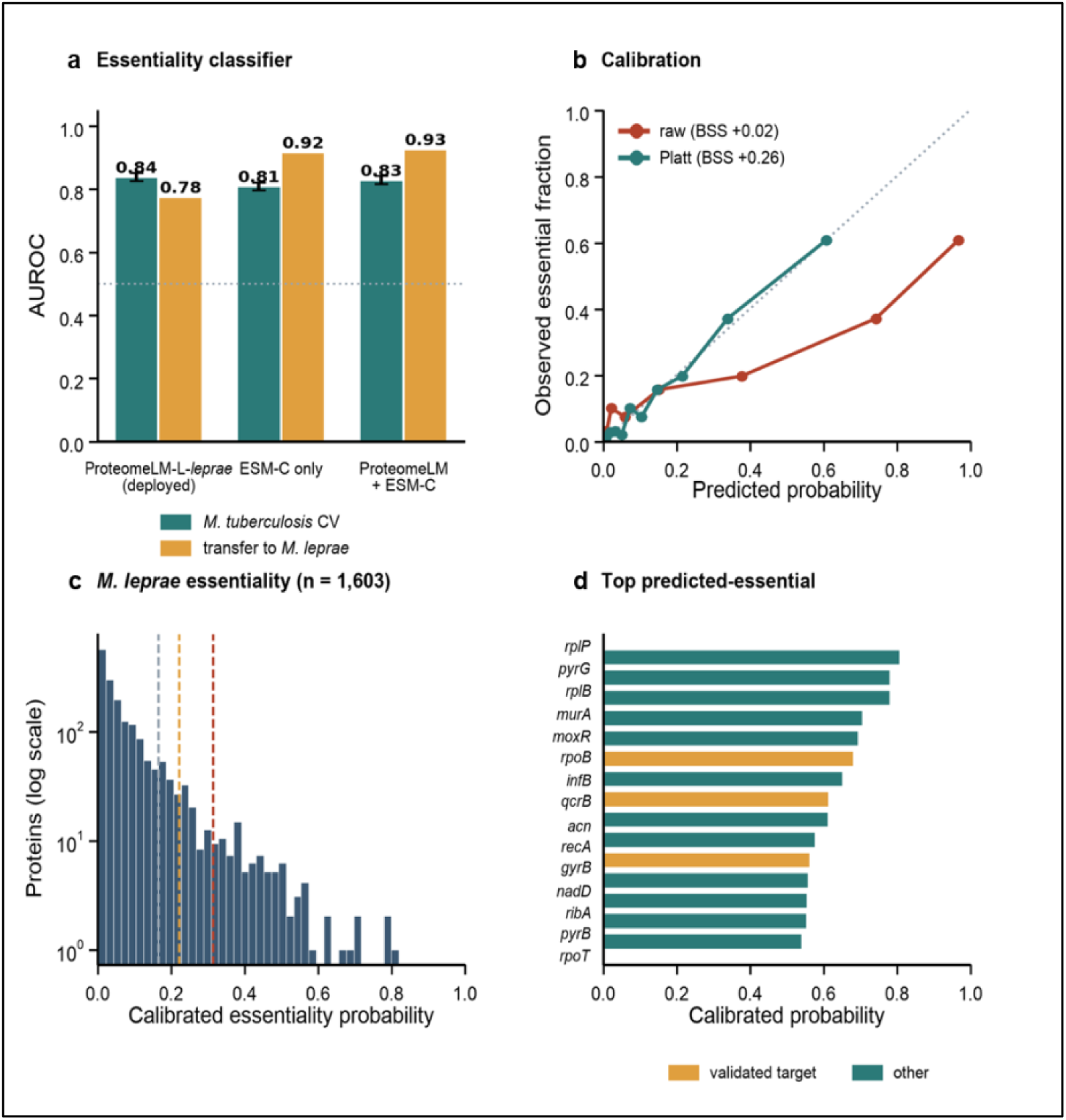
ProteomeLM-L-*leprae* cross-species essentiality prediction. (a) Homology-grouped five-fold cross-validation in *M. tuberculosis* (teal) and transfer to the orthologue-aligned *M. leprae* set (amber) for the three feature sets. Error bars show one standard deviation across folds. (b) Reliability of the predicted probabilities before and after Platt scaling, with Brier skill scores (BSS). (c) Distribution of calibrated ProteomeLM-L-*leprae* essentiality probabilities for the 1,603 *M. leprae* proteins (dashed lines indicate the class boundaries of 0.165, 0.222 and 0.315). (d) The 15 highest-scoring proteins; amber bars indicate validated anti-mycobacterial drug targets.

**Table 2.** Performance of the ProteomeLM-L-*leprae* essentiality classifier: The first three columns give homology-grouped cross-validation in *M. tuberculosis*, and the last column gives transfer of the classifier to *M. leprae*. Brier scores are for the uncalibrated probabilities. Essential genes make up 16.0 % of the labelled *M. tuberculosis* set, so a model that predicted that same proportion for every gene would reach a Brier score of 0.134. Measured against that reference, the three feature sets give Brier skill scores of +0.02, −0.15 and −0.05. Platt scaling of the deployed model raises its Brier skill score to +0.26 and leaves the ordering of the proteins unchanged. Each of the 241 *M. leprae* proteins carries the experimentally determined label of its *M. tuberculosis* orthologue, so a classifier that simply repeated that label would reach an AUROC of 1.000. The column therefore measures how well a feature set recovers sequence similarity rather than how well it predicts essentiality, and the choice of model rests on the cross-validation columns alone.

| Feature set / model | *M. tuberculosis*<br>CV AUROC | *M. tuberculosis*<br>CV AUPRC | Brier<br>score | Transfer<br>AUROC (*M. leprae*) |
| --- | --- | --- | --- | --- |
| ProteomeLM-L-*leprae*,<br>contextual features<br>(deployed) | $0.84 \pm 0.01$ | 0.54 | 0.132 | 0.78 |
| ESM-C only (baseline) | $0.81 \pm 0.01$ | 0.46 | 0.154 | 0.92 |
| ProteomeLM + ESM-C<br>ensemble | $0.83 \pm 0.01$ | 0.50 | 0.141 | 0.93 |

### 2.8 Database Statistics

The HANSEN Statistics page provides a quantitative overview of the structural and functional features available across the *M. leprae* proteome. It summarises the availability and composition of monomeric, oligomeric and ligand-containing structural models, together with associated feature layers across different prediction methods. Model quality is characterised using distributions of pLDDT scores and PAE, providing a more informative assessment. Comparative analyses further examine method-specific effect sizes, associations between functional categories and protein-level correlations. An integrated per-protein table combines structural model availability, oligomeric state, ligand information, model-confidence metrics, predicted binding pockets, B-cell epitope propensity, target-prioritisation scores and essentiality scores, enabling systematic evaluation of the breadth, quality and convergence of computational predictions across the proteome.

### 2.9 Benchmarking Model Accuracy and Confidence Measures

Across the proteome, the four modelling methods produced high-confidence models for most proteins (Fig. 4, Table S28). Comparing the monomeric models, Chai-1 and AlphaFold 3 gave the highest mean pLDDT at 85.6 and 85.3, with 90% and 89% of models reaching a mean pLDDT above 70. Boltz-1 and Boltz-2 were lower at 81.2 and 80.6. Including the oligomeric models raises the Boltz-2 mean to 82.0 but leaves the other three methods almost unchanged, because 704 of the 726 oligomeric models are registered to Boltz-2. Of those 704, 564 were co-folded by Boltz-2 and 140 were assembled by rigid-body superposition of Boltz-2 monomers onto template assemblies. Confidence ranking declined gradually for longer proteins, many of which are likely to contain multiple domains, consistent with expectations for *ab-initio* modelling.

**Fig. 4.**
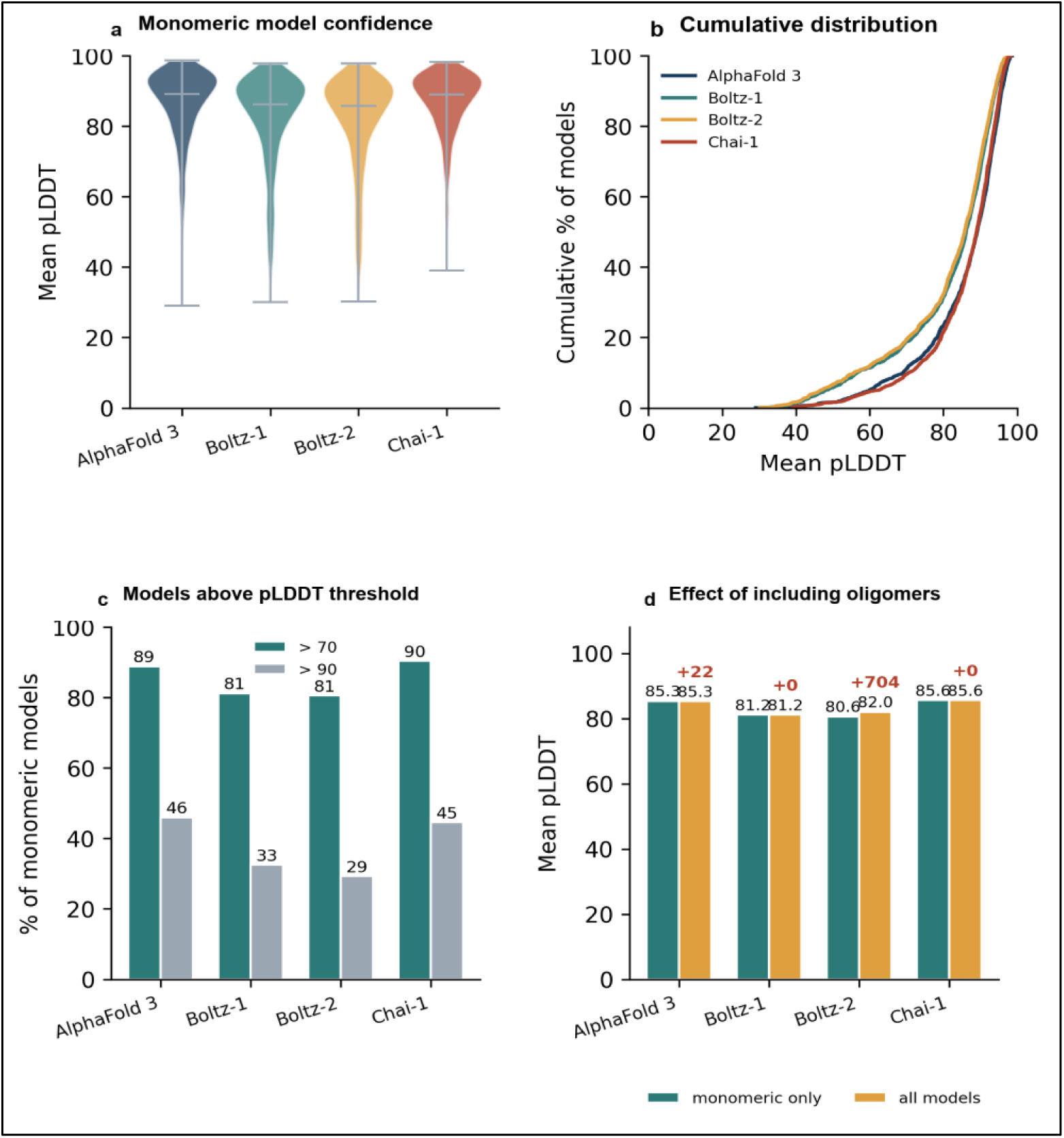
Proteome-wide model confidence. (a) Distribution of mean pLDDT by method for the monomeric models, shown as violin plots with medians. (b) Cumulative distribution of mean pLDDT, monomeric models. (c) Percentage of monomeric models with a mean pLDDT above 70 or above 90, by method. (d) Effect of including the oligomeric models on each method’s mean pLDDT, with the number of oligomeric models added shown in red. Mean pLDDT is the average of the per-residue pLDDT values over the Cα positions of each model.

For the seven proteins with an experimentally determined structure, agreement with the models was high(Fig. 5, Table S29). Mean TM-scores across the seven references were 0.90 for AlphaFold 3, 0.89 for Chai-1, 0.85 for Boltz-2 and 0.82 for Boltz-1, with a median Cα-RMSD of 1.9 Å. Six of the seven references were reproduced closely by all four methods, with TM-scores between 0.80 and 1.00. The exception was the Holliday junction branch migration protein RuvA (ML0482, 7OA5), where Boltz-1 and Boltz-2 returned TM-scores of 0.21 and 0.34 against AlphaFold 3 at 0.77 and Chai-1 at 0.73. RuvA carries a mobile C-terminal domain whose orientation differs between the monomeric models and the crystallised octamer, and excluding this case raises the Boltz-1 and Boltz-2 means to 0.92 and 0.93. Predicted pLDDT tracked the accuracy measured against the experimental structures, so it is a useful guide to local model reliability. Within each protein, the correlation between per-residue pLDDT and Cα-lDDT has a median of 0.76 and ranges from 0.32 to 0.90 across the 28 protein and method combinations (Fig. 5b). Pooling all residues gives 0.64, from 4,624 residue and method pairs, or 1,156 residues counted once per method (Fig. 5a).

**Fig. 5.**
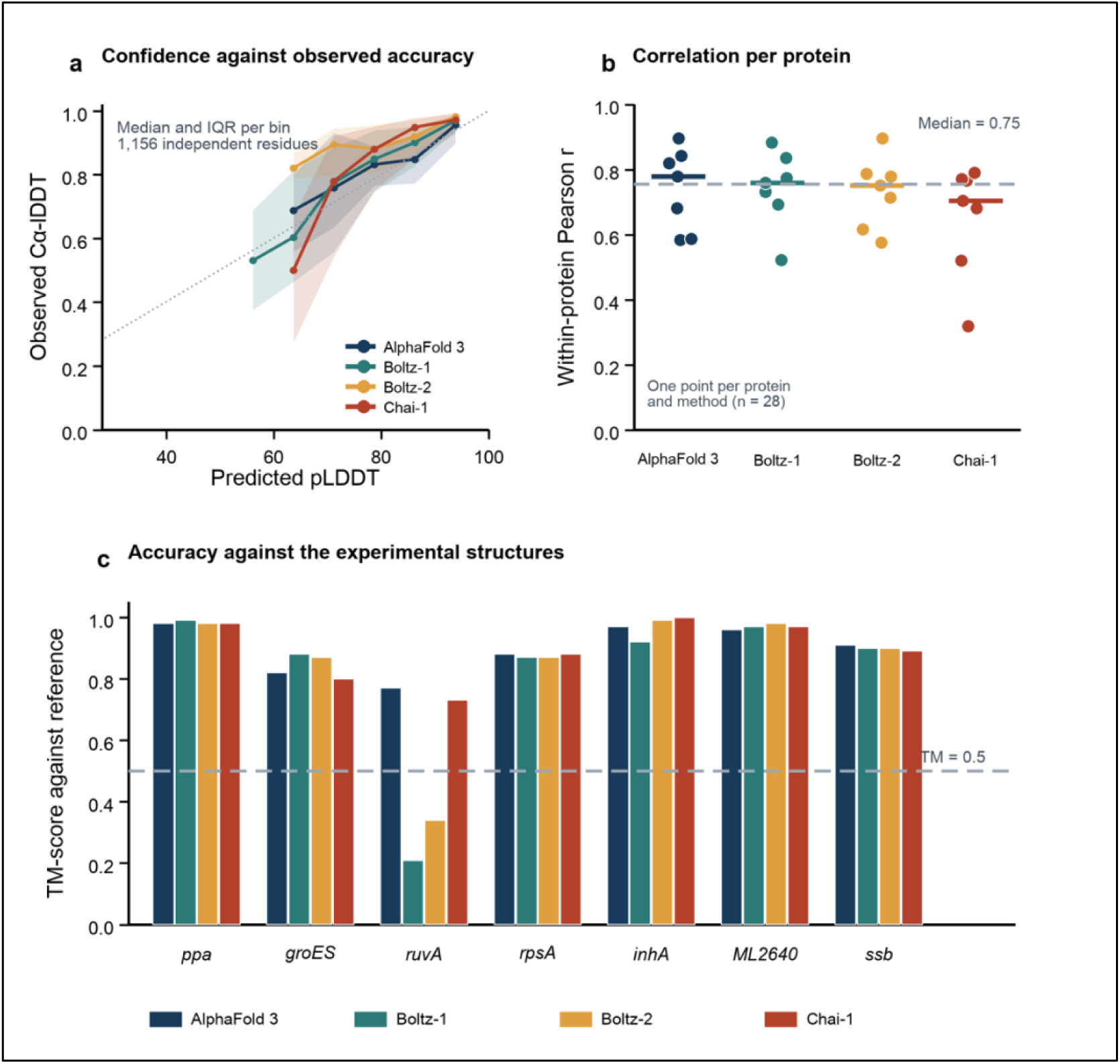
Accuracy against the *M. leprae* experimental structures. (a) Predicted pLDDT against observed Cα-lDDT, binned by method, showing the median and interquartile range within each bin; the standard-error bars of the previous version were computed over residues counted once per method and have been removed. The diagonal denotes perfect calibration. (b) Within-protein Pearson correlation between predicted pLDDT and observed Cα-lDDT, one point for each of the 28 protein-and-method combinations, with the median for each method marked. (c) TM-score for each predicted model against its experimental reference; the dashed line marks a TM-score of 0.5.

### 2.10 Cross-method concordance

Because experimental structures are available for only a few proteins, agreement between prediction methods provides the main internal measure of reliability for the remainder of the proteome. Among the proteins modelled by more than one method, pairwise TM-scores were generally high. Boltz-1 and Boltz-2 were the most concordant pair, with a mean TM-score of 0.83 and a per-residue pLDDT correlation of r = 0.96 (Fig. 6, Table S30). For each protein, we also identified the method pair with the highest TM-score. The median best-pair TM-score was 0.94, and 93% of proteins reached at least 0.5. Agreement increased with model confidence. Strong cross-method agreement is here defined as a best-pair TM-score of at least 0.5. The 115 proteins that do not meet it, 7.2% of the 1,599 proteins modelled by more than one method, were concentrated among models with low pLDDT and should be interpreted cautiously.

**Fig. 6.**
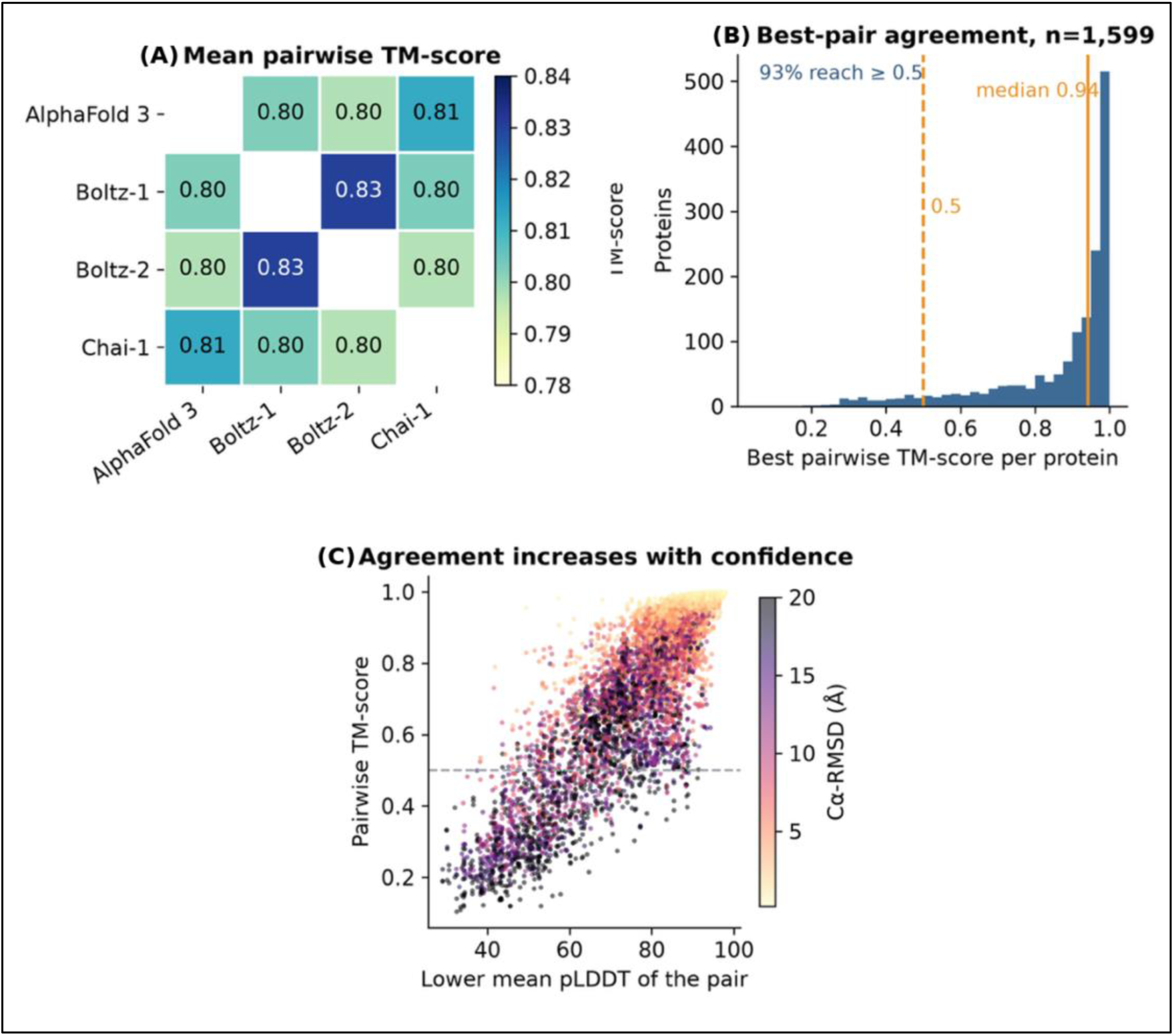
Cross-method concordance. (a) Mean pairwise TM-score between methods. (b) Distribution of the highest pairwise TM-score for each protein. (c) Pairwise TM-score against the lower mean pLDDT of the two models, coloured by Cα-RMSD, showing that cross-method agreement increases with model confidence. Values were derived from 9,582 pairwise comparisons across the proteins modelled by more than one method.

#### 2.10.1 Concordance with AFDB

The AlphaFold Protein Structure Database (AFDB) already provides a model for almost every *M. leprae* protein, so we asked how closely the HANSEN models reproduce it. We compared all four HANSEN methods with the AFDB model for 1,598 proteins, giving 6,379 model pairs. The two sets agree well. Half of the proteins scored above a TM-score of 0.934 for AlphaFold 3, 0.901 for Chai-1, 0.894 for Boltz-1 and 0.892 for Boltz-2, and between 86 and 90% of proteins reached the TM-score of 0.5 that indicates a shared fold. Agreement was higher still at the local level, where the median Cα-lDDT ranged from 0.948 to 0.971.

Where the scores were low, the models remained plausible. Slightly more than half of the low-scoring cases were proteins in which each domain was reproduced correctly but the domains were arranged differently, which a single superposition cannot capture. PknB is a clear example. Its whole-chain TM-score is only 0.44 to 0.49, yet its catalytic kinase domain matches the AFDB model to within 1.3 to 3.2 Å, and the difference lies in a disordered 55-residue linker that every method places 15 to 18 Å away. The same pattern appears in coiled-coil proteins such as Smc, in modular enzymes, in membrane-anchored proteins and in ribosomal proteins that only become ordered inside the assembled ribosome. Once the comparison is restricted to residues that both models regard as confident, more than 98% of proteins with a definable confident core reach a TM-score of 0.5 by every method. That denominator is 1,503, 1,435, 1,431 and 1,489 proteins for AlphaFold 3, Boltz-1, Boltz-2 and Chai-1 respectively, or 90 to 94% of the comparison set, because the excluded proteins are precisely those without at least 20 residues that both models regard as confident. Expressed over the full comparison set, the figures are 89 to 93%, and the sentence should be read with the denominator stated, which shows that the remaining disagreement sits in flexible and disordered regions.

Because the AFDB models are themselves predictions, this comparison measures reproducibility rather than accuracy. We therefore also asked whether AFDB stands apart from the four HANSEN methods by comparing all five sources with one another. Each source agrees with the other four to a similar degree, from 0.899 for Boltz-2 and Chai-1 to 0.910 for AlphaFold 3, with AFDB at 0.907, so AFDB is best regarded as a fifth member of the same ensemble rather than an independent standard. Agreement was highest for the proteins that matter most for drug discovery, averaging 0.922 to 0.962 across the 50 high-priority targets of the calibrated ranking. Two of those 50 targets (4%) have at least one method scoring below a TM-score of 0.7 against the AFDB, namely the riboflavin biosynthesis protein RibA and the DNA ligase LigA, and LigA falls at or below 0.51 for three of the four methods. No protein in the high-priority tier now falls below 0.5 for all four methods. The serine/threonine protein kinase PknB, moves to rank 84 in the strong-candidate tier once the essentiality probabilities were calibrated. LigA is one of the fifteen highest-scoring proteins for predicted essentiality and holds rank 13 of the calibrated ranking. These are differences in inter-domain packing between two sets of predictions rather than demonstrated errors, but they are exactly the proteins for which a single rigid-body superposition is uninformative, so a per-target agreement flag is reported in Table S20 and is being added to the prioritisation interface, and the tier mean is reported together with the count below threshold, and 139 of the 140 proteins on which all four methods disagreed with AFDB belong to the lowest-priority tier (Supplementary Note SN1).

HANSEN adds 726 oligomeric assemblies with interface confidence, template-derived ligands, pocket and ligandability predictions, residue-level epitope propensity, cross-species essentiality, an integrated target ranking, and the Lep-AMR module.

### 2.11 Recovery of Validated Drug Targets by the Priority Score

To test whether the Target Priority Score captures biologically meaningful targets, we assembled a reference set of 24 validated targets represented in the ranking. Validated targets were enriched near the top of the list. Five of the 24 were recovered among the top 50 candidates, compared with 3% expected by chance, 15 among the top 200 and 22 among the top 400, against random expectations of 12% and 25% respectively (Fig. 7, Table 3). The highest-ranked validated targets included DNA gyrase subunit B (*gyrB*, rank 3), cytochrome bc1 complex subunit QcrB (*qcrB*, rank 12), RNA polymerase subunit beta (*rpoB*, rank 6), thymidylate synthase ThyX (*thyX*, rank 32) and arabinosyltransferase C (*embC*, rank 44); these five are the validated targets that fall inside the high-priority tier. Some known targets ranked lower because the score integrates druggability, pocket and tractability score as well as prior validation. The ranking should therefore be interpreted as a triage tool, not a definitive ordering of biological importance.

**Fig. 7.**
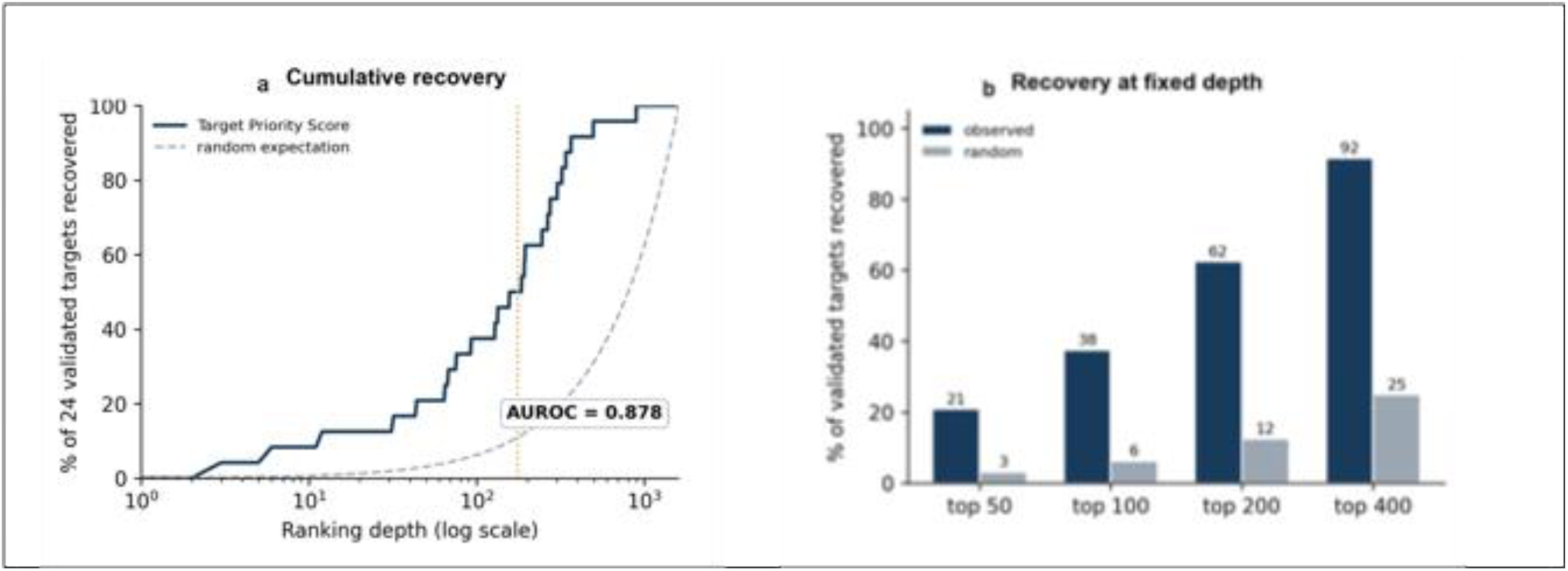
Recovery of validated drug targets by the Target Priority Score. (a) Cumulative percentage of the 24 validated reference targets recovered as a function of ranking depth, compared with random expectation; the dotted line marks the combined 175-protein high-priority and strong-candidate tiers. (b) Percentage of validated targets recovered at fixed ranking depths, compared with random expectation.

**Table 3.** Recovery of the 24 validated reference targets within top-ranked candidates, compared with the expectation under random ordering of the 1,603 ranked proteins.

| Ranked subset | Validated recovered | Recovered (%) | Random expectation (%) |
| --- | --- | --- | --- |
| Top 50 | 5 of 24 | 20.8 | 3.1 |
| Top 100 | 9 of 24 | 37.5 | 6.2 |
| Top 200 | 15 of 24 | 62.5 | 12.5 |
| Top 400 | 22 of 24 | 91.7 | 25.0 |
| Top 806 | 23 of 24 | 95.8 | 50.3 |

Across the 1,603 ranked proteins the 24 validated targets are recovered with an AUROC of 0.878 (95% bootstrap confidence interval 0.823 to 0.921) and an AUPRC of 0.105 against a baseline of 0.015. Enrichment is significant at every depth examined: 5 of 24 within the top 50 (6.7-fold over random, hypergeometric p = 6.6 × 10⁻⁴), 9 of 24 within the top 100 (6.0-fold, p = 6.0 × 10⁻⁶), 15 of 24 within the top 200 (5.0-fold, p = 8.0 × 10⁻⁹) and 22 of 24 within the top 400 (3.7-fold, p = 5.8 × 10⁻¹²). The lowest-ranked validated target is the ATP synthase subunit c, atpE (ML1140), at rank 895. It is a small, highly hydrophobic membrane subunit penalised by the size and tractability terms, which illustrates the limits of a score built for soluble enzymes. The composition of the validated set, with rank and score for each member, is given in Table S26. Removing the annotation contribution leaves the ranking almost unchanged (Spearman ρ = 0.995, top 50 Jaccard index 0.852), so lexical matching does not drive the recovery, but a residual circularity remains because the essentiality classifier and the validated target set both are derived from *M. tuberculosis*, in which drug targets are enriched for essential genes.

Removing each component in turn and repeating the benchmark shows that no single component performs as well as the full score, though the annotation term comes closest, with an AUROC of 0.866 against 0.878. The binding-site component is 1.0 for all 1,603 proteins because the consensus pocket score reaches its maximum for every protein, and a value that is the same everywhere cannot change the order of the ranking. This component accounts for 0.25 of the score directly and 0.045 through tractability, so 29.5% of the weight has no effect on the ranking, which is set by annotation, essentiality and model quality. The full component-by-component comparison is given in Supplementary Note SN3.

### 2.12 Implementation and functional evaluation of Lep-AMR Module

Lep-AMR module provides a browser-accessible workflow for detecting antimicrobial-resistance-associated variants in Oxford Nanopore amplicon data from *M. leprae*. The platform analyses the *rpoB*, *folP1* and *gyrA* resistance-determining regions individually or as a three-gene panel. When the target is unknown, it screens uploaded reads against the reference library and selects the best-supported amplicon. Detected variants are translated into amino-acid substitutions and reported with sequencing depth, filter status and antimicrobial category. HARP is an open database of predicted effects for 80,902 systematically generated missense substitutions on protein stability and molecular interactions across three *M. leprae* drug targets (14). Lep-AMR links eligible missense calls to HARP for structural interpretation. Run monitoring, EPI2ME reports, mutation tables, logs and complete outputs are available through the interface.

The example workflow was demonstrated with a single barcode *folP1* amplicon dataset that we recently acquired from India (Madhusmita Das et al., 2025, Unpublished). Lep-AMR correctly detected *folP1* and processed 2,306 reads comprising 656,409 bases, with a mean read length of 284.7 bp and a mean quality score of 11.7. Approximately 77% of reads mapped to the reference amplicon, producing a reported mean coverage of 1,697.6-fold. Variant analysis identified one C>T substitution at position 89 of the reference amplicon that passed the quality filters. No insertions or deletions were detected. Lep-AMR translated the call into the FolP1 P55L substitution in ML0224 and classified it as associated with dapsone resistance. This example demonstrates the ability of Lep-AMR to transform Nanopore amplicon reads into an interpretable gene- and drug-specific report.

## 3. DISCUSSION

HANSEN addresses a major structural knowledge gap in leprosy research by providing a unified structural and functional resource for the *M. leprae* proteome. *M. leprae* remains experimentally intractable and has far fewer experimentally determined structures than related mycobacteria. By integrating modern structure prediction, confidence metrics, pocket detection, epitope mapping, essentiality prediction and target prioritisation, HANSEN provides a practical means of accelerating hypothesis generation and structure-guided drug discovery.

A key strength of HANSEN is the integration of multiple features around each gene name. Researchers can search for a protein, inspect its structural models and confidence metrics, examine ligandability and predicted binding sites, view epitope propensity and assess prioritisation within one interface. This integration reduces reliance on separate tools and supports target discovery, functional annotation and structure-guided inhibitor design. Table S8 shows how the different features combine for individual proteins.

The inclusion of oligomeric assemblies is another important feature. Many proteins function as multimers, and monomeric structures alone may not reveal relevant interfaces, allosteric sites or ligand-binding sites. By representing oligomeric models, classifying assembly states and displaying oligomeric PAE, HANSEN provides a richer view of possible assemblies. Confidence and PAE JSON files, model mmCIF files, pocket outputs and epitope predictions are handled according to their method, assembly type and data format.

The four predictors reproduce the AFDB models closely across the proteome, and the AFDB models themselves remain within the predictor ensemble. The comparison measures reproducibility across independent implementations rather than accuracy. The differences that remain are informative. They fall almost entirely in regions that every method already flags as low confidence, and in the relative placement of domains in modular, coiled-coil and membrane-anchored proteins, where a single rigid-body superposition does not describe the molecule. For those proteins the individual domains are reproducible, and the inter-domain arrangement is not, and the models are best interpreted domain by domain in both resources. Where the four methods disagree with one another, they also disagree with the AFDB, so cross-method agreement can be used as an internal reliability assessment for the proteins that have no experimental structure.

The most informative extension available from the present data is to determine whether oligomerisation exposes pockets absent from the monomeric models. The pocket-detection pipeline and the oligomeric models are both already implemented, so the comparison requires no new modelling, and interface pockets identified on high-priority targets would constitute a new result rather than a re-description of existing models. A retrospective docking assessment against the template-derived ligands would convert the stated suitability for virtual screening into a measured outcome.

HANSEN is a prediction resource, and its outputs are hypotheses for experimental testing. Every model carries per-residue pLDDT and pairwise PAE. These quantify the internal consistency of the prediction and localise the regions in which it is self-consistent, but they do not measure agreement with the native structure. Accuracy is established here by comparison with the seven available experimental structures and by concordance across four independent predictors, and both assessments are limited by the small experimental set.

Five properties of the model set define the conditions under which it should be used, and each is recorded for every model. First, coordinates are reported as predicted, without geometry regularisation. Clashscore, Ramachandran and rotamer statistics are not computed, so stereochemical validation is required before docking or free-energy calculation. Second, template-derived ligands were retained where the closest protein-to-ligand heavy-atom distance lay between 1.2 and 6.0 Å. Hydrogen-bond donor-to-acceptor distances are 2.6 to 3.2 Å, so the lower bound of this window admits van der Waals overlap, and the ligand coordinates therefore define the binding site rather than a refined pose. Third, monomer models are apo. Apo prediction of a metalloenzyme yields an active site that is not organised around the absent ion, so cavity volumes and druggability scores computed on those sites are not comparable with holoenzymes. This applies to high-priority metalloenzyme targets including ribA, glmS and the folate enzymes, for which the corresponding oligomeric models, which carry template-derived cofactors, should be used for active-site analysis. Fourth, a single conformer is retained per protein, so cryptic and allosteric sites detectable only across a conformational ensemble are not represented. Fifth, oligomeric stoichiometry is inherited from the PDB biological-assembly record, which is PISA-derived and not independently verified, and hetero-oligomeric partners are assigned by best sequence match within the *M. leprae* proteome. The template identifier and construction method are recorded for every assembly in Table S2, so both can be checked.

To our knowledge, HANSEN is the first proteome-wide leprosy-focused resource to integrate *M. leprae* genome-derived annotations with proteome-wide structural, functional, essentiality and therapeutic-target features. Leproma established an important genomic foundation by providing access to the reduced TN genome, coding sequences and extensive pseudogene repertoire (*28*). MycoBrowser later incorporated *M. leprae* into a comparative mycobacterial genomics interface for gene-level annotation and cross-species comparison (*29*). HARP supports antimicrobial-resistance interpretation by reporting predicted structural effects of systematic missense substitutions in RNA polymerase, dihydropteroate synthase and DNA gyrase (*14*). HANSEN complements these resources by connecting genomic locus tags and sequences to predicted monomeric and oligomeric structures, binding-site predictions, essentiality estimates and target-prioritisation features. Its Lep-AMR component links Nanopore amplicon data to nucleotide and amino-acid substitutions and drug-specific resistance interpretation.

In conclusion, HANSEN is a comprehensive web-based structural and functional resource for the *M. leprae* proteome. It addresses a longstanding barrier by providing broad access to predicted structures, model-confidence metrics, PAE visualisations, oligomeric assemblies, ligands, predicted binding pockets, B-cell epitope propensity, essentiality and AI-assisted target prioritisation. By linking structural models with functional and translational annotations through UniProt accession and gene names, HANSEN supports drug-target discovery, binding-site analysis, epitope mapping and systems-level interpretation. Future developments may incorporate validated biochemical data, resistance-associated mutations, comparative pathogen–host specificity metrics, expanded ligand-docking outputs, updated essentiality datasets and additional immune-epitope predictions. These additions would further strengthen HANSEN as a platform for translational research in leprosy and related neglected tropical diseases.

## 4. MATERIALS AND METHODS

### 4.1 Proteome Dataset and Protein Annotation

The *M. leprae* protein dataset was assembled from UniProt records and incorporated into the HANSEN relational database. Each protein record was keyed by its gene name in UniProt (*20*) and linked to available protein descriptions, functional annotations, pathway information and cross-references. Users can query the database by UniProt accession, gene name, protein name, functional term, ligand or amino-acid sequence.

### 4.2 Structural Modelling Strategy

HANSEN uses complementary structure-prediction methods. Structural modelling was performed with Seq3D, an in-house automated workflow for proteome-scale structure prediction (Rees et al., manuscript in preparation). A sequence-similarity search against PDB records identified candidate homologues. All proteins were first modelled as monomers. Where a suitable template was identified, its annotated biological assembly guided the stoichiometry of an additional homomeric model. Using AlphaFold 3, we modelled 193 such homo-oligomers and with Boltz2, we modelled homo- and hetero-oligomers, of which 506 homomeric and 198 heteromeric records were retained. With the 22 AlphaFold 3 homomers that have no Boltz-2 counterpart this gives 528 homo-oligomeric and 198 hetero-oligomeric records, or 726 in total, which is the composition reported in Table 1.

At the time of analysis, ABCFold ran AlphaFold 3, Boltz-1 and Chai-1 within one workflow (*18*). Boltz-2 models were generated separately after its release and retained as an additional prediction set for comparison (*19*). For each model, HANSEN stores assembly type, prediction method, structure-file path, method-specific ranking score and rank. Database-linked model records were used for summary statistics. Confidence metrics are provided for model interpretation.

### 4.3 Oligomeric Assembly Modelling

HANSEN includes predicted homo- and hetero-oligomeric assemblies generated with template-informed AlphaFold 3 and Boltz-2 workflows. Candidate templates were identified by complementary searches of PDB-derived libraries. Foldseek compared modelled *M. leprae* monomer structures with PDB templates (*30*), and sequence level local *blastp* compared the corresponding amino-acid sequences with PDB chain sequences (*31*). Candidate templates were then ranked by a combined score that favoured high identity, query coverage, sequence and structural similarity, and agreement with the PDB biological assembly (oligomeric state and chain stoichiometry). The top-ranked templates were selected. For hetero-oligomers, each *M. leprae* query protein was matched by Foldseek to a chain within a multi-chain PDB assembly, and the highest-scoring assembly was used as the template. The template’s PDB biological assembly defined the global stoichiometry of the model complex. Each protein entity in the template was then assigned to its top *blastp* hit in the *M. leprae* proteome, with all chains of that entity inheriting the assignment, producing an explicit map between template chains and *M. leprae* partners. For homomeric input files in Boltz2, the YAML files contain repeated protein sequence according to the template stoichiometry, whereas for heteromeric input files, the full partner protein composition was encoded.

For complete-chain assemblies with incorrect relative placement, an automated script was used to treat each predicted chain as a rigid body and superposed it on the corresponding template chain with sequence-aligned Cα atoms and a Kabsch fit (*32*). This changed the relative placement of chains without altering their internal conformation. Corrected assemblies were verified for chain count, centroid placement, overlap, completeness and ligand clashes.

Where both workflows produced an assembly for the same protein, a single representative was retained. Of the 193 AlphaFold 3 homomeric assemblies, 171 had a Boltz-2 counterpart and are represented by the Boltz-2 model, and the remaining 22 were retained as AlphaFold 3 assemblies. The database therefore holds 726 oligomeric model records: 564 co-folded by Boltz-2, 140 assembled by rigid-body superposition of Boltz-2 monomers onto template assemblies, and 22 from AlphaFold 3. These 726 records resolve to 660 distinct model files and 546 distinct template assemblies; the 140 superposed records derive from 46 distinct template assemblies, of which PDB 8V9K, the ribosome, accounts for 53. The distinct assemblies, records, template PDB identifiers, stoichiometry, construction method and interface confidence is provided in the Table S2. The platform distinguishes monomers from oligomers and classifies assemblies by chain count, enabling comparison of confidence across oligomeric states.

### 4.4 Model confidence: pLDDT and PAE

Where available, per-residue pLDDT values were read from model mmCIF records and summarised at residue and model level (*33*). PAE matrices were read from method-specific JSON files. Each PAE value estimates the expected positional error for residue *i* when the predicted and reference structures are aligned on residue *j*. Lower values indicate greater confidence in relative placement(*34*).

HANSEN stores and parses method-specific confidence files, including monomer PAE matrices, Boltz-2 monomer confidence files, AlphaFold 3 homomer PAE matrices and Boltz-2 oligomer PAE matrices. In oligomeric matrices, diagonal blocks describe within-chain confidence and off-diagonal blocks describe confidence in inter-chain placement. The interface predicted aligned error reported for each assembly is the mean of these off-diagonal values over pairs of residues on different chains whose representative atoms, Cβ or Cα where no Cβ is present, lie within 8 Å of one another. Per-residue pLDDT was read from the B-factor field of each model mmCIF, averaged over the Cα atoms of each chain to give the model mean. Residues were binned into the conventional bands of very high (above 90), confident (70 to 90), low (50 to 70) and very low (50 or below), which define the Mol* colouring and the residue-band counts stored with each model. Model-level confidence is summarised as the percentage of models with a mean pLDDT above 70 and above 90, and the percentage of residues above 70 (Table S28). No pLDDT cut-off was applied to model inclusion; thresholds are interpretive, and the model-quality term of the Target Priority Score uses mean pLDDT on a continuous scale. Observed local accuracy was measured as Cα-lDDT, computed without superposition over Cα pairs within a 15 Å inclusion radius at distance tolerances of 0.5, 1.0, 2.0 and 4.0 Å. For comparisons of predicted confidence with observed accuracy, a confident core was defined as residues at which both structures reach a pLDDT of 70 or above, requiring at least 20 such residues. Large PAE matrices are loaded on demand after the Mol* (*26*) structure to preserve interface responsiveness.

### 4.5 Ligand and cofactor annotation

Ligand annotations were obtained from model metadata and HETATM records in structural mmCIF files. Exclusions were based on component identity rather than Foldseek or BLAST scores. Standard amino-acid and water molecules were excluded. For template-derived ligands added to oligomers, components were retained only when the closest heavy-atom distance to the protein was between 1.2 and 6.0Å. The lower bound of this window is permissive: a separation of 1.2Å is a steric clash, and template-derived ligand placements should be inspected before use rather than assumed to be physically valid. Valid ligand codes are displayed with the relevant models and linked to external chemical resources.

For Boltz-2 oligomeric modelling, ligand SMILES strings from the best-matching templates were supplied in the YAML input files. This enabled selected assemblies to be modelled with template-informed ligand representations. The AlphaFold 3 oligomeric models deposited in HANSEN do not currently contain ligand coordinates.

### 4.6 Pocket detection and druggability

HANSEN integrates three complementary pocket-prediction approaches. AF2BIND (25) provides residue-level binding propensity, while P2Rank (*24*) and fpocket (*23*) identify putative ligand-binding pockets from structural features. Predicted pockets are displayed in a panel linked to Mol*, allowing users to focus on pocket residues and assess candidate sites alongside structural confidence and functional annotation. Pocket characteristics also contribute to target prioritisation. AF2BIND operates on AlphaFold 2 pair representations. The reported 21,204 candidate binding-site residues across 1,603 proteins are the top-ranked positions retained for each model.

### 4.7 Sequence, feature and functional annotation

HANSEN links structural models to protein sequence and functional annotation, integrating domains, motifs, gene and protein names, functional descriptions, cross-references and pathway associations. This allows a predicted pocket to be interpreted in relation to catalytic residues, conserved motifs, protein-family annotation, essentiality and pathway role.

### 4.8 B-cell epitope propensity mapping

B-cell epitope propensity was computed with DiscoTope-3.0 (*27*) and treated as a residue-level characteristic rather than a protein-level estimate. Scores for supported AlphaFold 3 and Boltz-2 models are displayed with residue number, residue type, threshold information and Mol* focus functions. DiscoTope-3.0 uses spatial context and can therefore identify discontinuous candidate epitopes in predicted structures.

### 4.9 ProteomeLM-derived Contextual Essentiality

Gene essentiality in *M. leprae* was predicted with ProteomeLM-L-*leprae* by cross-species transfer from *M. tuberculosis*. The *M. tuberculosis* H37Rv proteome and the 1,603 HANSEN proteins were encoded with ESM-C and passed through ProteomeLM-L to yield whole-proteome contextual embeddings. Because ProteomeLM processes at most 512 proteins per pass, each proteome was embedded in overlapping, genome-ordered windows, and the per-protein embeddings were averaged across windows. ProteomeLM is distributed as a masked language model with a protein–protein interaction module.

A logistic-regression essentiality classifier was trained on the *M. tuberculosis* contextual embeddings using the saturating-transposon labels of *DeJesus et al*. (*35*). Five-fold cross-validation was grouped by MMseqs2 clusters at a sequence identity of at least 40%, so that sequence-similar proteins did not span the training and test folds (*36*). Under homology-grouped cross-validation the model reached a mean Area Under the Receiver Operating Characteristic (AUROC) of 0.84. Applying it across species to the 241 *M. leprae* proteins that align *to M. tuberculosis* orthologues gave a transfer AUROC of 0.78. Only 39 of those 241 proteins carry an essential label, and those labels are the DeJesus classification of the *M. tuberculosis* orthologue rather than an experimental measurement in *M. leprae*. Assigning each protein the classification of its orthologue therefore scores an AUROC of 1.000 by construction, so the transfer figure measures recovery of sequence similarity and cannot be used to compare models. The homology-grouped cross-validation (Fig. 3a) is the only usable measure of performance. The calibrated essentiality probabilities for *M. leprae* appear on the essentiality page and provide the main ProteomeLM-L-*leprae* term in the HANSEN Target Priority Score.

Of the 1,603 proteins, 1,408 (87.8%) have an identifiable *M. tuberculosis* orthologue, which is consistent with earlier comparative analyses of the two genomes. The figure of 241 quoted above is not the orthologue count but the subset of those orthologues that carries a saturating-transposon essentiality label and can therefore be used for transfer evaluation. Of these, 39 are labelled essential and 202 non-essential. The transfer AUROC is computed on that labelled subset alone.

### 4.10 Target prioritisation

*M. leprae* proteins were ranked for structure-guided drug discovery and experimental follow-up using a weighted Target Priority Score. The score combines five components, each scaled from 0 to 1: the ProteomeLM contextual essentiality score; a binding-site score, taken as the greater of the P2Rank/fpocket consensus-pocket score and the AF2BIND score; a functional-annotation score; a Boltz-2 model-quality score; and a tractability score.

On a 0–100 scale,

*Target Priority Score = 100 × [(0.35 × essentiality) + (0.25 × binding-site) + (0.20 × annotation) + (0.10 × model quality) + (0.10 × tractability)]*.

For the annotation score, the functional characteristics contribute as follows: an Enzyme Commission (EC) number, 0.16; a metabolic-pathway assignment, 0.16; a catalytic-activity statement, 0.12; an annotated active or binding site, 0.12; and a Gene Ontology assignment, 0.08. A further 0.045 is added for each match to a predefined set of 36 terms associated with essential mycobacterial processes; this lexical contribution is capped at 0.40, and the annotation score at 1.0. The 36 terms are listed verbatim in the Table S27. Because these terms can name target families, the benchmark of Section 2.11 is repeated with the lexical contribution set to zero and both results are reported, so that recovery of validated targets is not measured by string matching.

Tractability score: This is computed as (0.45 × binding-site) + (0.30 × size-suitability) + (0.25 × enzymatic indicator). The size-suitability term is 1.0 for chains of 120–750 residues, 0.75 for 751–1,200, 0.65 below 120, 0.45 above 1,200, and 0.60 where length is unavailable. The enzymatic indicator is 1.0 for proteins carrying an EC number or an enzyme or binding-related annotation term for example, an enzyme class such as kinase, ligase or hydrolase, or the terms active site, binding site or cofactor, and 0.45 otherwise. Binding-site features contribute to both the explicit binding-site score and the tractability score by design.

Proteins are assigned to priority tiers by their Target Priority Score alone, with no other criterion: high-priority (≥67), strong candidate (≥61), moderate candidate (≥50) and exploratory (<50). The ranking is a computational shortlist for target discovery, not a validated measure of essentiality, biological indispensability or druggability. High-ranking candidates therefore require expert biological review, assessment of pathway relevance, comparison with established mycobacterial drug targets and experimental validation before progression into drug-discovery workflows.

### 4.11 Lep-AMR – an interface for processing Nanopore AMR Data

Lep-AMR was implemented within HANSEN as a local workflow for analysing Oxford Nanopore MinION amplicon-sequencing data from *M. leprae*. The reference library covers the principal resistance-determining regions of *rpoB*, *folP1* and *gyrA*, associated with rifampicin, dapsone and fluoroquinolone resistance, respectively. Users upload a compressed directory of FASTQ or FASTQ.GZ files from one barcode and may select one target, analyse the full three-gene panel or allow automatic target detection. For automatic detection, sampled reads are compared with the three reference amplicons and the best-supported target is selected.

Validated reads, a sample sheet and the selected reference FASTA are passed to the EPI2ME Nextflow wf-amplicon workflow, version 1.2.2 (*37*). Reads are filtered with a minimum length of 80 bp, a minimum mean quality score of 10, at least 40 reads and variant-calling coverage of at least 20. Alignment uses minimap2 version 2.28 (*38*), and haploid small-variant calling uses Medaka version 2.2.0 with the dna_r10.4.1_e8.2_400bps_hac@v4.2.0 model. Resulting VCF calls are translated into nucleotide and amino-acid substitutions. Each substitution is assigned to its gene name by matching the translated amplicon against the corresponding protein sequence. The interface reports the gene name residue-level substitution, sequencing depth and filter status and provides a HARP link for missense variants. It does not currently project variants automatically onto a HANSEN three-dimensional model as same models exist in HARP for the three known drug targets in *M. leprae*. Uploaded data and generated outputs are deleted automatically after 24 hours or may be deleted manually.

### 4.12 Web implementation

HANSEN was implemented as a Flask/Jinja2 web application backed by a relational database. The platform provides protein-centric query pages, interactive structure-visualisation pages, statistics dashboards, target-prioritisation and essentiality pages, a GO browser, help documentation and contact information. Each protein page integrates identifiers, functional annotation, sequence features, cross-references, monomeric and oligomeric structure models, model-quality metrics, pocket predictions, AF2BIND binding-site predictions, ligand annotations, epitope predictions and interactive Mol* visualisation.

### 4.13 Quality Assessment and Benchmarking

To evaluate the reliability, internal consistency and translational utility of the HANSEN structural atlas, five complementary benchmarking analyses were performed as follows:

1. Residue and model-level confidence distributions were compared across monomeric structures generated by AlphaFold 3, Boltz-1, Boltz-2 and Chai-1.
2. Cross-method structural concordance was assessed for proteins modelled by more than one method. Shared residues were sequence-matched, Cα atoms were superposed with the Kabsch algorithm and pairwise similarity was calculated using TM-score (*39*) and Cα-RMSD. TM-score was computed with the standard iterative fragment-seed search rather than from a single global superposition. Per-residue confidence concordance was assessed from correlations between matched pLDDT values.
3. Accuracy against experimental structures was evaluated for proteins with exact experimental structural matches. Predicted models were superposed onto the corresponding experimental structures, and structural agreement was quantified using TM-score, Cα-RMSD and per-residue Cα-lDDT. Calibration of predicted confidence was assessed by comparing model-derived pLDDT values with observed Cα-lDDT values, thereby estimating whether high-confidence predicted regions corresponded to regions of higher experimental structural agreement.
4. The integrated Target Priority Score was benchmarked against a reference set of validated or biologically supported targets. The set included leprosy MDT targets and *M. leprae* orthologues of established mycobacterial drug targets. Receiver operating characteristic analysis and descriptive top-k early recovery were used to test whether known targets were enriched near the top of the ranking. AUROC and AUPRC with bootstrap confidence intervals, and a hypergeometric p-value at each ranking depth, are reported in Section 2.11. The composition of the validated reference set is given in full in the Table S26.
5. The HANSEN monomers were compared with the corresponding models in the AFDB. The AFDB proteome release for *M. leprae* TN (UP000000806_272631_MYCLE, model version v6) was paired one-to-one with the HANSEN monomer set by UniProt accession, giving 1,598 proteins and 6,379 model pairs. Residue correspondence was established from UniProt numbering without structural alignment, and the 84 proteins whose modelled construct boundaries differ between the two sets were aligned globally and compared over the aligned region. Each pair was scored by TM-score normalised by the AFDB chain length, GDT-TS(*40*), Cα-RMSD after Kabsch superposition and superposition-free Cα-lDDT, together with per-residue pLDDT concordance and Cα-based secondary-structure agreement. All ten pairwise combinations of the five model sources were computed on the same scale so that the AFDB models could be placed within the predictor ensemble. The TM-score implementation was verified against the reference TM-align program on 400 randomly chosen pairs, giving a Pearson correlation of 0.996 to 0.998 and a median difference of zero. Full methods, tables and figures are given in Supplementary Note SN1.

Structure parsing and coordinate extraction were performed with Gemmi (*41*). Statistical analyses, performance metrics and classification benchmarks were implemented with scikit-learn (*42*).

## Supporting information

Supplementary Workbook 1

Supplementary Workbook 2

## ACKNOWLEDGMENTS

Author contributions

S.C.V. and S.M. conceived the study. S.C.V. designed the HANSEN resource and relational database schema, supervised data integration, and wrote the first draft of the manuscript. S.M. and R.R. contributed to structural modelling, feature integration, data analysis, and manuscript revision. A.M., M.M., A.A., C.B., K.S., and M.D. contributed to data analysis and manuscript writing. T.L.B. and R.A.F. provided supervision and resources and critically revised the manuscript. S.C.V and R.R. contributed equally to this work. All authors read and approved the final manuscript.

## Data availability

HANSEN is available at https://hansen-leprosy.medschl.cam.ac.uk/. The site provides interactive structure visualisation, proteome statistics, and target-prioritisation and essentiality browsers, and the complete set of 7,122 models covering 1,603 proteins can be downloaded per protein or in bulk in TSV, CSV, mmCIF and JSON format, so that every analysis reported here can be repeated on the underlying data. The resource is provided for non-commercial research and education. Its components carry different licences. Boltz-1 and Boltz-2 are MIT, Chai-1 and ProteomeLM are Apache 2.0, DiscoTope-3.0 is under an academic licence, and the AlphaFold 3 models are subject to the AlphaFold 3 Model Parameters Terms of Use, with bulk redistribution of the AlphaFold 3-derived coordinates confirmed as compliant with those terms. All content is computational prediction intended for hypothesis generation and must not be used to make, support or inform clinical or diagnostic decisions.

The code and metric implementations underlying Supplementary Materials are available at https://github.com/sundeep-vedithi/HANSEN.

## Artificial Intelligence Use

AI-assisted tools were used for English grammar and usage under author supervision. The authors reviewed and approved all scientific content, analyses and conclusions.

## SUPPLEMENTARY MATERIALS

**Supplementary Note SN1:** Structural concordance between the AlphaFold Protein Structure Database and the HANSEN monomeric models, covering 1,598 proteins and 6,379 model pairs, with Figs. S1 to S6.

**Supplementary Note SN2:** Cross-species essentiality transfer, model selection and calibration.

**Supplementary Note SN3:** Additional methodological notes. Figs. S7 and S8 show the protein page and the target-prioritisation browser of the web interface. Tables S28 to S30 give proteome-wide model confidence by modelling method, accuracy against the seven experimental reference structures, and cross-method concordance.

**Supplementary Workbook 1 (Tables S1 to S8, S26 and S27):** All registered structure models ranked by confidence (S1); all oligomeric assembly models with inferred template and confidence (S2); ligand and cofactor records for the oligomeric assemblies (S3); and all 1,603 proteins ranked by pocket features (S4), predicted B-cell epitope propensity (S5), Target Priority Score (S6) and predicted essentiality (S7), together with the integrated per-protein table (S8).

**Supplementary Workbook 2 (Tables S9 to S25):** Comparison coverage and construct differences (S9, S10); global agreement, confidence concordance and the confident core (S11 to S13); agreement by chain length, secondary structure and predicted localisation (S14, S15, S21); discordant proteins, the split between local fold and domain packing, and named case studies (S16, S17, S24); five-way concordance among the five model sources (S18); agreement by target-priority tier and the 50 top-ranked targets individually (S19, S20); per-protein results for all 6,379 comparisons (S22); validation of the TM-score implementation (S23); and the domain-resolved comparison for PknB (S25).

Table S26 gives the 24 validated reference targets with gene, ML identifier, UniProt accession, priority rank, score and tier. Table S27 lists verbatim the 36 terms used by the lexical component of the annotation score.

