## Supplementary Material for "HANSEN: An Integrated Structural and Functional Proteome Resource for Structure-Guided Drug Discovery in *Mycobacterium leprae*"

**Supplementary Materials:**

### Supplementary Note SN1: Structural concordance between the AlphaFold Protein Structure Database (AFDB) and HANSEN monomer models

#### 1. Scope

We compared the AlphaFold Protein Structure Database (AFDB) proteome release for *Mycobacterium leprae* TN with the monomer models in HANSEN. The AFDB release UP000000806_272631_MYCLE uses model version 6 and contains 1,602 single-chain models. The HANSEN monomer models were generated with four co-folding predictors, namely AlphaFold 3, Boltz-1, Boltz-2 and Chai-1. The analysis covers 1,598 proteins and 6,379 model pairs.

We limited the comparison to monomers because the AFDB proteome release contains only single-chain models. HANSEN contains 726 oligomeric model records that have no counterparts in this release. These comprise 564 records co-folded with Boltz-2, 140 reconstructed by superposing monomeric models onto a template assembly and 22 generated with AlphaFold 3. We therefore excluded these records.

#### 2. Model sets and residue correspondence

Of the 1,602 AFDB entries, 1,598 correspond uniquely to *M. leprae* loci for which HANSEN monomers are available. The other four are secondary UniProt accessions linked to locus labels already used by other entries. They are O05755 at ML2219, Q9CD20 at ML2569, Q7AQ19 at ML1911 and P0A5D0 at ML2428A, which is a locus in its own right rather than a second accession for ML2428. In each case, we retained the accession whose UniProt sequence exactly matches the modelled chain.

One HANSEN protein has no corresponding AFDB entry. ML1191 (Q7AQ85, fatty acid synthase) is 3,076 residues long and exceeds the length limit of the AFDB proteome release. Four other loci, ML1180, ML1181, ML1183 and ML2692, have monomer models but no UniProt accessions. No AFDB models can therefore be assigned to them. Of the 1,603 loci with HANSEN monomers, 1,598 consequently have an AFDB reference (Table S9).

The comparison includes 1,598 pairs for AlphaFold 3, 1,595 for Boltz-1, 1,595 for Boltz-2 and 1,591 for Chai-1. These differences arise from the composition of the HANSEN monomer set rather than from the comparison procedure.

We established residue correspondence without structural alignment. Of the 6,379 pairs, 3,325 have identical modelled sequences. Another 2,717 differ at one position and were matched using UniProt residue numbering. The remaining 337 pairs required a global Needleman-Wunsch alignment because the boundaries of their modelled constructs differ. Every one-residue difference occurs at position 1. UniProt records the initiator residue as methionine, whereas the genomic translations used to generate the HANSEN models retain the valine or leucine encoded by the GTG and TTG start codons of *M. leprae*. No other positions differ within these pairs, so residue correspondence is unaffected.

Eighty-four proteins have different modelled chain lengths in the two model sets (Table S10). The four largest differences occur in ML0842, which has 611 residues in AFDB and 411 in HANSEN, ML1985, which has 606 and 798 residues respectively, ML0015, which has 232 and 47 residues respectively, and ML0021, which has 155 and 340 residues respectively. These differences arise from gene boundary annotation rather than modelling. Each comparison was performed over the aligned region.

### 3. Quality Metrics

We calculated all metrics using Cα atoms from matched residue pairs and treated the AFDB model as the reference. TM-score (39) used the standard iterative fragment seed search and was normalised by AFDB chain length. We used the sequence-dependent form because both structures represent the same sequence. GDT-TS used cut-offs of 1, 2, 4 and 8 Å, while GDT-HA (43) used 0.5, 1, 2 and 4 Å. We evaluated lDDT (44), the superposition-free local distance difference test described by Mariani and colleagues, with Cα atoms, a 15 Å inclusion radius and the four standard tolerance thresholds. Global RMSD was calculated across all matched residues after one Kabsch superposition. Core RMSD was limited to residues within 5 Å of the reference after TM superposition, so it represents the shared substructure. We assigned secondary structure from Cα geometry with P-SEA (45) and compared it using Q3, a measure of three-state agreement. Confidence was based on the per-residue pLDDT values stored in each model's B-factor field.

We validated TM-score against TM-align using 400 randomly selected pairs that covered all four methods (Table S23). The correlation exceeded 0.996 for every method. The median difference was zero, and our implementation scored 0.0051 lower on average. TM-align always produced the higher score because it optimises a sequence-independent alignment. For models of the same sequence, the sequence-dependent score used here is more appropriate.

TM-score is normalised by AFDB chain length, so a HANSEN model covering only part of the reference cannot score much higher than its coverage fraction. Chain coverage is below 95% in only 15 or 16 comparisons per method. Removing these comparisons changes the median TM-score by no more than 0.002, so the proteome-wide values are unaffected. ML0842, a cysteine desulfurase with 611 residues in AFDB and 411 in HANSEN, illustrates this effect. Its TM-score ranges from 0.66 to 0.67, close to the coverage ratio of 0.67. However, the 411 shared residues superpose with an RMSD of 0.8 to 1.5 Å and an lDDT of 0.986 to 0.991. We excluded these comparisons from the classification in Section 6.

### 4. Global agreement

The four predictors closely reproduce the AFDB models across the proteome (Table S11 and Fig. S1). Across the four methods, the median TM-score relative to AFDB ranges from 0.892 to 0.934. The proportion of proteins that meet or exceed the TM-score threshold of 0.5 for a shared fold range from 86.4 to 90.3%. Local agreement is higher than global agreement for every method, with median lDDT-Cα values ranging from 0.948 to 0.971.

**Table S11. Global agreement with the AFDB reference***

| Method | n | Median TM-score | TM-score IQR | Median GDT-TS | Median lDDT | Median RMSD (Å) | Median core RMSD (Å) | TM ≥ 0.5 | TM ≥ 0.9 |
|---|---|---|---|---|---|---|---|---|---|
| AlphaFold 3 | 1,598 | 0.934 | 0.755–0.984 | 0.902 | 0.971 | 2.93 | 0.85 | 90.3% | 59.6% |
| Boltz-1 | 1,595 | 0.894 | 0.677–0.964 | 0.843 | 0.948 | 5.54 | 0.99 | 86.8% | 48.8% |
| Boltz-2 | 1,595 | 0.892 | 0.683–0.964 | 0.839 | 0.948 | 6.09 | 0.99 | 86.4% | 47.7% |
| Chai-1 | 1,591 | 0.901 | 0.700–0.970 | 0.853 | 0.950 | 4.79 | 0.98 | 87.8% | 50.1% |

**Selected columns. The complete table is provided in worksheet S11 of the accompanying workbook.*

The TM-score distributions are unimodal at high values and have a low-value tail rather than a broad spread. For AlphaFold 3, the mean of 0.839 is lower than the median of 0.934. Mean whole-chain RMSD is between eight and ten times greater than the median RMSD over the shared substructure. Whole-chain RMSD is therefore a poor summary of proteome-wide agreement because it is dominated by the multi-domain proteins analyzed in Section 6.

AlphaFold 3 is closest to the AFDB reference, as expected from the shared architecture of the two systems. Among the 1,591 proteins compared with all four methods, AlphaFold 3 produces the highest TM-score for 1,165 proteins. The corresponding counts are 218 for Chai-1, 108 for Boltz-1 and 100 for Boltz-2. The median TM-score differs by 0.042 between AlphaFold 3 and Boltz-2, whereas the three non-AlphaFold predictors differ from one another by no more than 0.008.

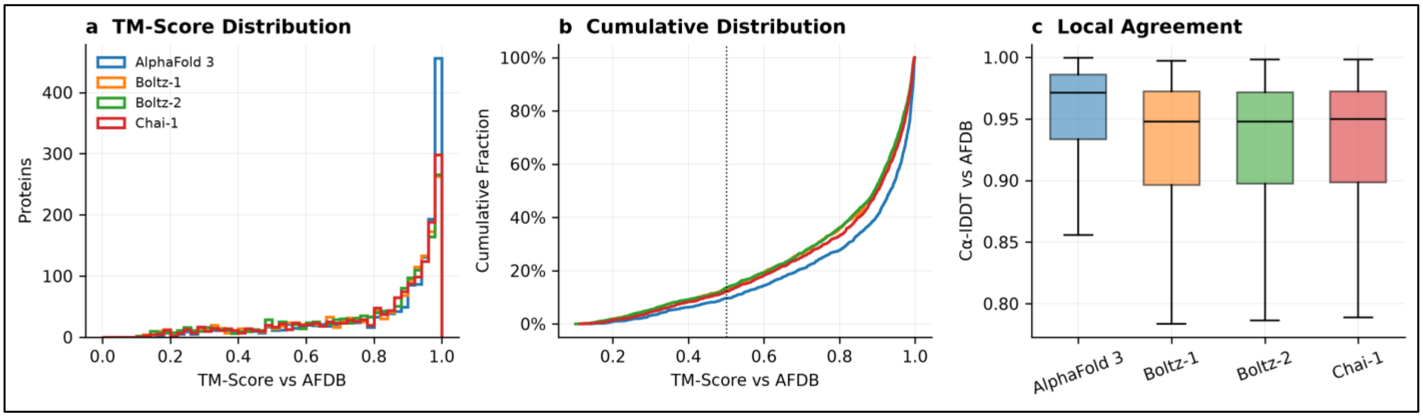


**Fig. S1.** Agreement between the AFDB models and the four HANSEN predictors across 6,379 model pairs. (a) Histograms show TM-scores relative to AFDB in 50 bins from 0 to 1, with one outline for each method. All four distributions have a single mode above 0.95 and a low-scoring tail. Table S11 reports their medians. (b) Cumulative distributions show the same data. The dotted vertical line marks 0.5, the conventional threshold for a shared fold. The proportions below this threshold are 9.7% for AlphaFold 3, 13.2% for Boltz-1, 13.6% for Boltz-2 and 12.2% for Chai-1. (c) Box plots show lDDT-Cα, a superposition-free measure of local agreement. The boxes cover the interquartile range; the line marks the median and the whiskers extend to 1.5 times the interquartile range. Outliers are omitted. The low-scoring tail of the TM-score distribution has no equivalent in the lDDT distribution because TM-score is sensitive to the relative placement of domains, whereas lDDT is not.

### 5. Model Confidence

All confidence values in this section were calculated for the 1,598 proteins compared here. They therefore differ slightly from the proteome-wide values in Table S28, which include 1,603 proteins. AlphaFold 3 and Chai-1 closely match the AFDB proteome mean of 85.7, whereas Boltz-1 and Boltz-2 are lower by 4.5 and 5.1 units respectively (Table S12 and Fig. S2c). Coordinate agreement does not follow this order. The median lDDT differs by only 0.002 among Boltz-1, Boltz-2 and Chai-1 (Table S11). The offset therefore reflects differences in the confidence estimates rather than in the coordinates.

**Table S12. Concordance of model confidence**

| Method | Mean AFDB pLDDT | Mean model pLDDT | Δ | Median pLDDT profile r | r (model pLDDT, TM) | r (AFDB pLDDT, TM) | ρ (min. pLDDT, TM) |
|---|---|---|---|---|---|---|---|
| AlphaFold 3 | 85.7 | 85.4 | −0.28 | 0.926 | 0.850 | 0.837 | 0.856 |
| Boltz-1 | 85.7 | 81.2 | −4.47 | 0.897 | 0.886 | 0.828 | 0.880 |

| | | | | | | | |
|---|---|---|---|---|---|---|---|
| Boltz-2 | 85.7 | 80.6 | −5.08 | 0.898 | 0.894 | 0.834 | 0.892 |
| Chai-1 | 85.7 | 85.6 | −0.03 | 0.906 | 0.829 | 0.850 | 0.865 |

Per-residue confidence profiles are consistent across the methods. The median within-protein Pearson correlation between the AFDB and model pLDDT profiles ranges from 0.897 to 0.926. This indicates that AFDB and the four predictors identify the same reliable regions in each chain.

Per-residue deviation is closely related to confidence (Fig. S2a). Across the 534,329 AlphaFold 3 residues, the median Cα deviation from AFDB is 0.34 Å when the lower of the two pLDDT values is at least 90, a group containing 329,784 residues. The deviation is 0.94 Å when this value is between 70 and 90, based on 122,978 residues, and 2.6 Å when it is between 50 and 70, based on 38,947 residues. It rises to 17.9 Å when the value is below 50, based on 42,620 residues. The deviation is therefore 52 times greater in the lowest-confidence bin than in the highest-confidence bin.

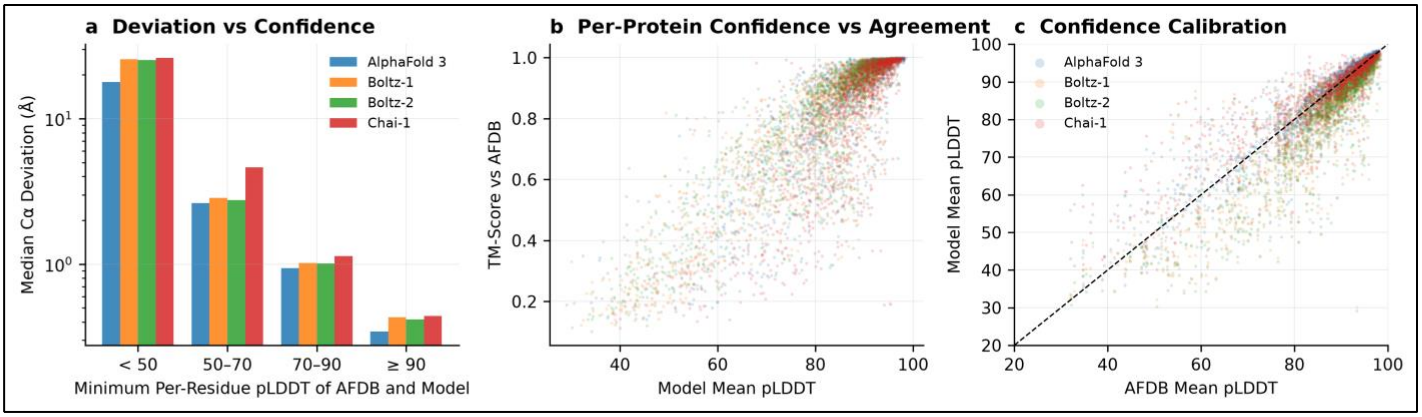


**Fig. S2.** Model confidence and structural agreement. (a) Median Cα deviation from AFDB after TM superposition, calculated per residue and grouped by the lower of the two pLDDT values. Bars are coloured by method and the vertical axis is logarithmic. The text reports the AlphaFold 3 values, and the other methods show the same pattern. (b) Mean model pLDDT is plotted against TM-score for each protein and method. The correlations are 0.850 for AlphaFold 3, 0.886 for Boltz-1, 0.894 for Boltz-2 and 0.829 for Chai-1. (c) Mean pLDDT for AFDB is plotted against each predictor. The dashed identity line shows that AlphaFold 3 and Chai-1 are closely calibrated to AFDB, while Boltz-1 and Boltz-2 lie below it (Table S12).

### 6. Low-confidence regions, local fold and domain packing

Limiting each comparison to residues with pLDDT values of at least 70 in both structures increases the mean TM-score by 0.067 to 0.083 and the mean lDDT by 0.031 to 0.042, depending on the method (Table S13 and Fig. S3b). Within this confident core, between 98.1 and 99.4% of proteins achieve a TM-score of at least 0.5.

lDDT does not require superposition, whereas TM-score does. Together, they distinguish two types of structural difference (Table S17). High lDDT with low TM-score indicates that the individual domains are reproduced but placed differently. Low values for both measures indicate a difference in local structure. For each method, 177 to 228 comparisons meet the first definition, while 149 to 192 score low on both measures. The first group contains longer proteins, with mean lengths of 339 to 392 residues compared with the proteome median of 283. Their mean whole-chain RMSD values are 13 to 17 times their mean core RMSD values, indicating rigid-body displacement.

**Table S17. Local fold and domain packing**

| Method | n | Same fold and packing (lDDT≥0.8, TM≥0.7) | Domain rearrangement (lDDT≥0.8, TM<0.7) | Divergent local structure (lDDT<0.8, TM<0.7) | Mean RMSD, rearranged (Å) | Mean core RMSD, rearranged (Å) | Mean rearranged length |
|---|---|---|---|---|---|---|---|
| AlphaFold 3 | 1,582 | 1,244 (78.6%) | 177 (11.2%) | 149 (9.4%) | 19.2 | 1.47 | 339 |
| Boltz-1 | 1,579 | 1,151 (72.9%) | 226 (14.3%) | 190 (12.0%) | 22.8 | 1.38 | 392 |
| Boltz-2 | 1,580 | 1,148 (72.7%) | 228 (14.4%) | 192 (12.2%) | 22.7 | 1.38 | 392 |
| Chai-1 | 1,575 | 1,171 (74.3%) | 215 (13.7%) | 178 (11.3%) | 20.7 | 1.50 | 365 |

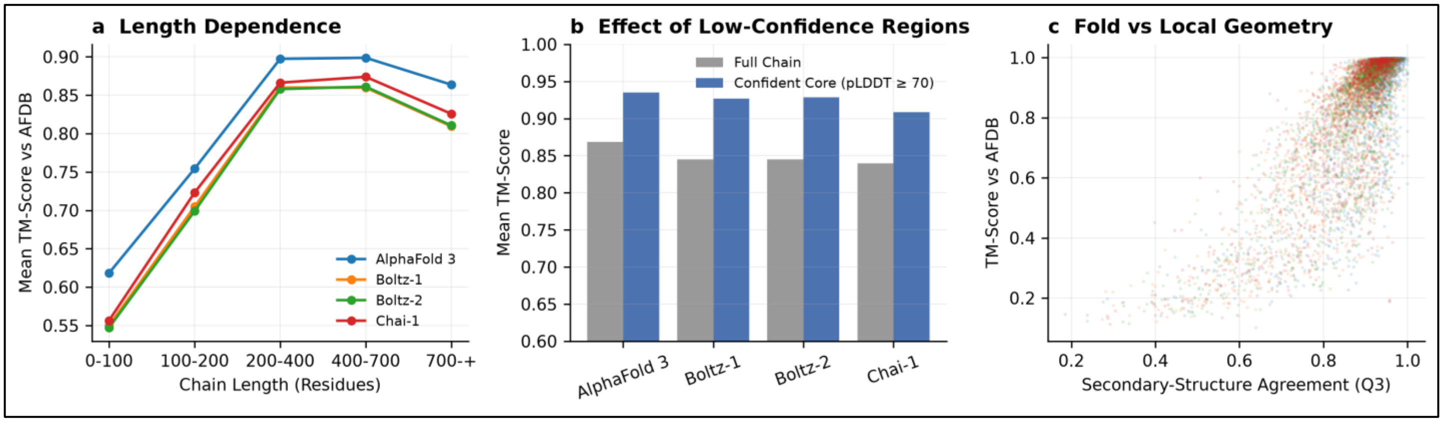


**Fig. S3.** Chain length, low-confidence regions and secondary structure. (a) Mean TM-score relative to AFDB is plotted by chain length, with one line for each method. For all four methods, the value rises from the shortest group to a maximum in the groups containing 200 to 400 and 400 to 700 residues, then falls in the longest group. The values are reported in Table S14. (b) Mean TM-score for the full chain is shown in grey alongside the same measure for the confident core in blue. The confident core comprises residues with pLDDT values of at least 70 in both the AFDB and HANSEN models. The increase for each method is reported in Table S13. (c) Three-state secondary structure agreement, Q3, is plotted on the horizontal axis against TM-score on the vertical axis. Each protein is represented by one point for each method. Secondary structure was assigned from Cα geometry with the P-SEA method. For AlphaFold 3, the mean Q3 is 0.946 among proteins with a TM-score of at least 0.9 and 0.728 among the 155 proteins with a TM-score below 0.5. The corresponding value across the complete set is 0.906. The correlation between Q3 and TM-score ranges from $r = 0.78$ to 0.80 across the four methods. Proteins that disagree globally therefore also differ in secondary structure across part of the chain. This is consistent with the absence of regular structure in the low-confidence segments described in Section 5.

In total, 45 proteins show this domain rearrangement signature across all four methods and have a confident AFDB model. The criteria are a mean lDDT of at least 0.85, TM-scores below 0.70 for all four methods and a mean AFDB pLDDT of at least 80 (Table S24). Of these proteins, 24 fall into four groups defined by their UniProt annotations. Two are elongated coiled-coil proteins. These are Smc (ML1629, 1,203 residues, TM 0.32 to 0.37) and ScpB (ML1369, 231 residues, TM 0.39 to 0.48). Twelve are modular enzymes or multi-domain proteins joined by flexible linkers. They are the phthiocerol polyketide synthase subunit (ML2353, 1,489 residues, TM 0.60 to 0.66), the non-ribosomal peptide synthase Nrp (ML1996, 1,401 residues, TM 0.63 to 0.64), the protein kinase PknB (ML0016, 622 residues, TM 0.44 to 0.49), RecA (ML0987, 711 residues, TM 0.53 to 0.65), the penicillin-binding protein PonA (ML2688, 708 residues, TM 0.63 to 0.67), UvrC (ML0562, 647 residues, TM 0.38 to 0.56), the replication initiator DnaA (ML0001, 502 residues, TM 0.52 to 0.67), a DNA polymerase (ML0603, 371 residues, TM 0.57 to 0.69), an AAA+ ATPase (ML0510, 473 residues, TM 0.62 to 0.68), a RecF-like AAA protein (ML1120, 873 residues, TM 0.37 to 0.69), the dihydrolipoamide acetyltransferase E2 subunit (ML0861, 530 residues, TM 0.47 to 0.61) and the dehydrogenase SerA (ML1692, 528 residues, TM 0.62 to 0.65). Five are membrane-anchored proteins whose soluble domains are attached to one transmembrane segment. These are the Rieske subunit QcrA (ML0880, 394 residues, TM 0.51 to 0.62), the cytochrome oxidase subunit CtaF (ML0876, 139 residues, TM 0.46 to 0.69), a band-7 protein (ML1802, 374 residues, TM 0.49 to 0.62), FtsQ (ML0916, 341 residues, TM 0.60 to 0.69) and MmpS4 (ML2377, 154 residues, TM 0.60 to 0.63). Five adopt a defined conformation only within a larger assembly. These are the ribosomal proteins uL22 (ML1858, 175 residues, TM 0.60 to 0.67), uS17 (ML1854, 126 residues, TM 0.62 to 0.65) and bL32 (ML0173, 57 residues, TM 0.53 to 0.64), the nucleoid-associated protein Lsr2 (ML0234, 112 residues, TM 0.48 to 0.56) and the cell wall scaffold Wag31 (ML0922, 266 residues, TM 0.44 to 0.55).

### 7. Domain-resolved comparison for PknB

PknB (ML0016, P54744) is a serine/threonine protein kinase that ranks 84th, within the strong-candidate tier of the calibrated ranking, and has a TM-score below 0.5 for all four methods. We therefore compared its individual domains using the UniProt domain annotations (Table S25 and Fig. S4). Across the full chain, the TM-score ranges from 0.44 to 0.49, RMSD ranges from 25 to 45 Å and lDDT ranges from 0.92 to 0.96.

**Table S25. Domain-resolved comparison for PknB (ML0016) ***

| Segment | Residues | AFDB pLDDT | TM AlphaFold 3 | TM Boltz-1 | TM Boltz-2 | TM Chai-1 |
|---|---|---|---|---|---|---|
| Kinase domain (11–273) | 263 | 86.6 | 0.97 | 0.86 | 0.94 | 0.89 |
| Juxtamembrane linker (274–328) | 55 | 53.4 | 0.34 | 0.32 | 0.25 | 0.34 |
| Transmembrane helix (329–349) | 21 | 65.1 | 0.58 | 0.63 | 0.69 | 0.62 |
| PASTA 1–4 (352–622) | 271 | 88.6 | 0.86 | 0.72 | 0.70 | 0.67 |
| Whole chain (1–622) | 622 | 83.6 | 0.49 | 0.44 | 0.46 | 0.46 |

**Selected columns. The complete table is provided in worksheet S25 of the accompanying workbook.*

The catalytic kinase domain (residues 11 to 273) reproduces the AFDB model with TM-scores of 0.86 to 0.97 and RMSDs of 1.3 to 3.2 Å. The four extracellular PASTA repeats (residues 352 to 622) have TM-scores of 0.67 to 0.86. AFDB models the 55-residue juxtamembrane linker (residues 274 to 328) with low confidence, and all four methods displace it by 15 to 18 Å from the reference (Table S25). The whole-chain TM-score therefore reflects this one low-confidence linker rather than the kinase domain.

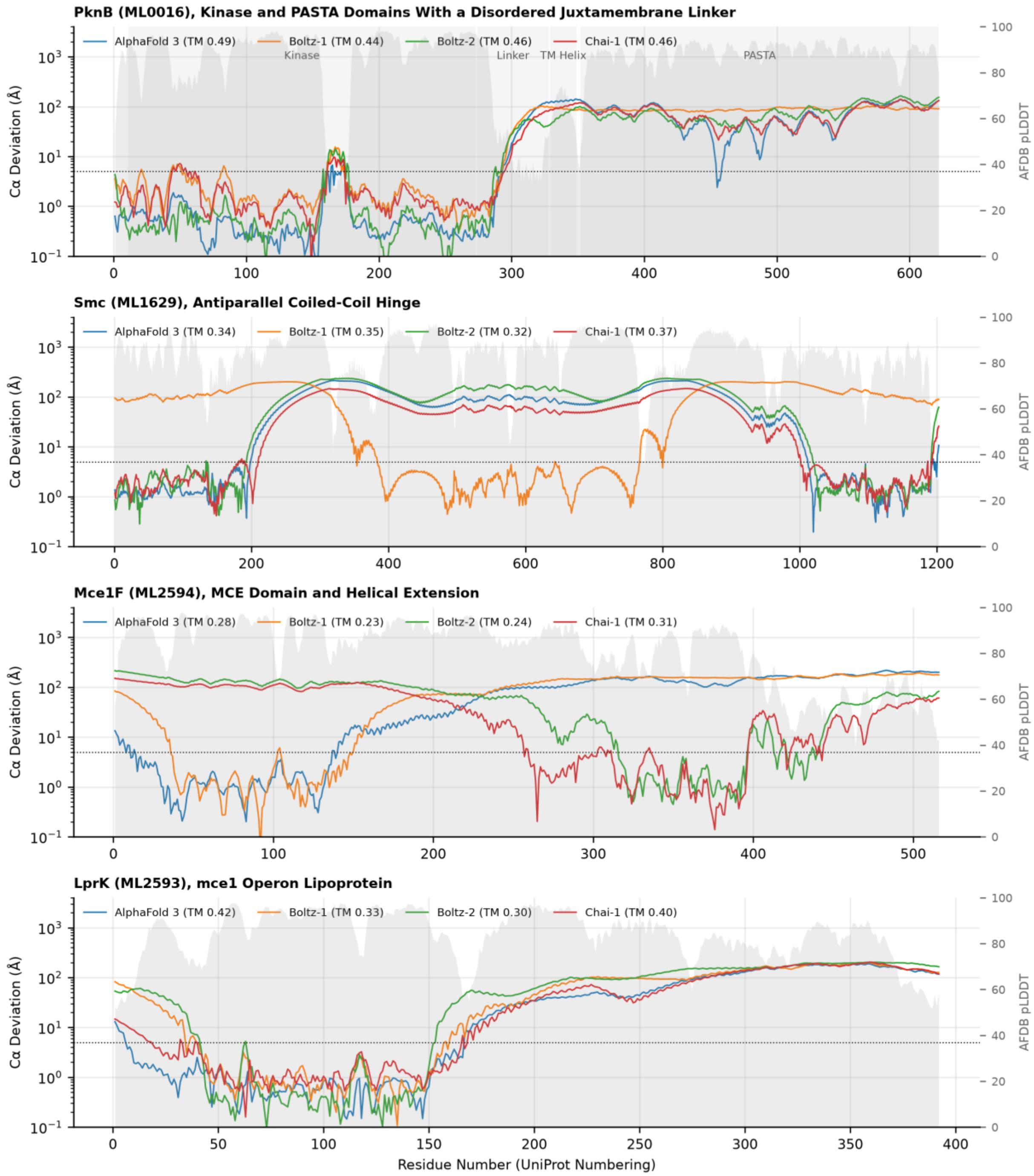


**Fig. S4.** Per-residue Cα deviation from AFDB after TM superposition for four proteins showing two types of disagreement. Each panel shows UniProt residue number horizontally and Cα deviation from 0.1 to 4,000 Å on the logarithmic left axis. The dotted line marks 5 Å. Grey shading shows AFDB pLDDT on the right axis. Each coloured line represents one method, whose TM-score is given in the legend. In PknB (ML0016), the UniProt domains are shaded and labelled. Deviation remains close to 1 Å across the kinase domain, which covers residues 11 to 273, before increasing by two orders of magnitude at the juxtamembrane linker, which covers residues 274 to 328. In Smc

(ML1629), each method follows low deviation across a different part of the chain, as expected when an antiparallel coiled-coil arm moves around a hinge. In Mce1F (ML2594) and LprK (ML2593), deviation remains close to 1 Å across the N-terminal MCE domain and rises steadily through the C-terminal helical extension, where AFDB pLDDT remains between 80 and 90. An abrupt step indicates a rigid-body difference. None of the panels shows the high deviation throughout the chain that would indicate a different fold.

### 8. Chain length, localisation and secondary structure

Agreement increases with chain length up to 700 residues (Table S14 and Fig. S3a). For AlphaFold 3, mean TM-score rises from 0.618 among the 143 proteins shorter than 100 residues to 0.899 among proteins with 400 to 700 residues. It then falls to 0.864 among the 104 proteins longer than 700 residues. The other methods follow the same pattern. TM-score normalisation is more stringent for short chains, which are also modelled with lower confidence. Their mean AFDB pLDDT is 72.7, compared with 85.7 across the complete set.

The 310 proteins with an annotated transmembrane segment show less agreement with AFDB than the 1,243 proteins with neither a transmembrane segment nor a signal peptide. The difference in mean TM-score is 0.090 for AlphaFold 3 and ranges from 0.097 to 0.115 for the other three methods (Table S21). All five model sources predict these protein structures without a membrane.

Secondary structure is reproduced consistently across the four methods (Table S15). Mean three-state agreement with AFDB is 0.906 for AlphaFold 3 and ranges from 0.861 to 0.862 for the other methods. Helices account for 46.3% of AFDB models and 46.9 to 48.9% of the predictor models. Strands account for 34.1% of AFDB models and 33.6 to 35.1% of the predictor models. The largest difference is 2.6 percentage points, with no corresponding exchange between helices and strands. The median ratio of each model's radius of gyration to that of the reference is 1.001 for AlphaFold 3, 0.974 for Boltz-1, 0.965 for Boltz-2 and 0.987 for Chai-1. Values below one show that the models are up to 3.5% more compact, consistent with the domain packing differences in Section 6. P-SEA uses Cα geometry and assigns regular secondary structure more freely than a hydrogen bond method such as DSSP. Helix and strand content should therefore be used to compare model sets rather than as absolute values.

### 9. Discordant proteins

A total of 288 proteins meet at least one of two discordance criteria. They either have a TM-score below 0.5 for one method relative to AFDB or show a range of more than 0.3 TM-score units across the four methods (Table S16). Of these proteins, 140 have a TM-score below 0.5 for all four methods. Their median chain length is 120 residues, compared with the proteome median of 283 residues, and 66% are shorter than 150 residues. The median of their mean AFDB pLDDT values is 59.9, compared with the proteome median of 89.7. In addition, 79% have a mean AFDB pLDDT below 70.

We defined a method-specific failure as a protein for which one method produces a TM-score below 0.5 while the other three do not. The counts are 4 for AlphaFold 3, 9 for Boltz-1, 13 for Boltz-2 and 12 for Chai-1. Table S24 lists seven cases in which the other methods achieve at least 0.70. The four Chai-1 cases are the ESAT-6-like antigen EsxB (ML0050, 100 residues, TM 0.43 to 0.92), alanyl-tRNA synthetase AlaS (ML0512, 908 residues, TM 0.48 to 0.84), the RNA helicase RhlE (ML0811, 544 residues, TM 0.43 to 0.77) and a MerR-type regulator (ML2075, 251 residues, TM 0.46 to 0.85). The three AlphaFold 3 cases are ML1793 (101 residues, TM 0.29 to 0.89), a DUF3349 protein (ML2258, 100 residues, TM 0.34 to 0.91) and ML1430 (108 residues, TM 0.47 to 0.82). These are all short chains of 100 to 108 residues. Their mean lDDT values range from 0.705 to 0.970, showing that the local structure is retained and the difference lies in the relative placement of chain regions. Three proteins disagree with a confident AFDB model across more than half of their chain. They are Mce1F (ML2594, 516 residues, TM 0.23 to 0.31), LprK (ML2593, 392 residues, TM 0.30 to 0.42) and ML0293 (69 residues, TM 0.33 to 0.40). Mce1F and LprK are encoded in the mce1 lipid import operon. Their mean AFDB pLDDT values are 80.9, 83.3 and 83.7 respectively, while the corresponding model means are 61.3, 65.9 and 64.9. Per-residue profiles locate the differences (Fig. S4). The N-terminal MCE domain of LprK, residues 1 to 150, superposes to within 1 Å in all four methods. Deviation then rises through the C-terminal helical extension to more than 100 Å at residue 350. Mce1F shows the same profile, as does the rearranged protein Mce1B (ML2590), which has a transition near residue 200. The region that remains superposed differs among methods, as expected for a hinge motion rather than an incorrect local fold.

#### 10. Position of the AFDB models within the ensemble

We calculated all ten pairwise comparisons among the five model sources on the same scale (Table S18 and Fig. S5). AFDB and AlphaFold 3 are the most concordant pair, with a median TM-score of 0.938. For each source, we averaged its median TM-scores against the other four. The five means span only 0.011. By this measure, AFDB ranks second behind AlphaFold 3 and lies within the ensemble.

**Table S18. Five-way concordance (median TM-score)**

| Method | AlphaFold DB (v6) | AlphaFold 3 | Boltz-1 | Boltz-2 | Chai-1 | Mean vs others |
|---|---|---|---|---|---|---|
| AlphaFold DB (v6) | 1.000 | 0.938 | 0.895 | 0.894 | 0.903 | 0.907 |
| AlphaFold 3 | 0.938 | 1.000 | 0.902 | 0.895 | 0.907 | 0.910 |
| Boltz-1 | 0.895 | 0.902 | 1.000 | 0.917 | 0.896 | 0.902 |
| Boltz-2 | 0.894 | 0.895 | 0.917 | 1.000 | 0.891 | 0.899 |
| Chai-1 | 0.903 | 0.907 | 0.896 | 0.891 | 1.000 | 0.899 |

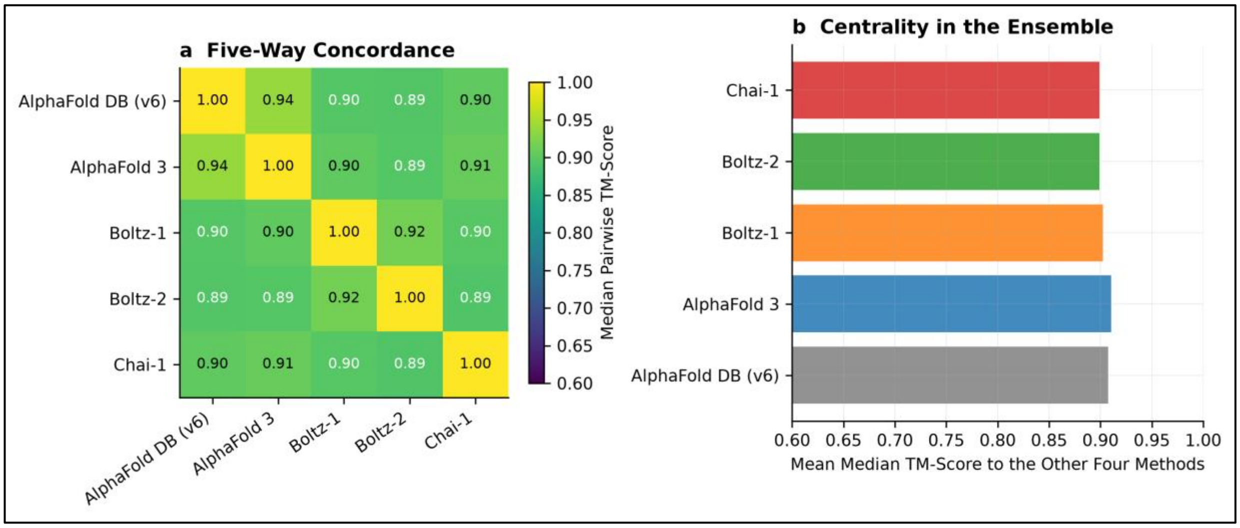


**Fig. S5.** Position of the AFDB models within the five-model ensemble. (a) The matrix shows the median pairwise TM-score for all ten unordered pairs of the five model sources. Values were calculated with the same code and normalisation used throughout this note. Cells are coloured from 0.60 to 1.00 and show the median value. The highest value outside the diagonal is 0.938 for the AFDB and AlphaFold 3 pair, while the lowest is 0.891. (b) Each bar is the mean of the four values outside the diagonal in the corresponding row of panel (a). It therefore represents the mean of the median TM-scores for one source against the other four. By this measure, the AFDB models rank second among the five sources behind AlphaFold 3.

Agreement among the four HANSEN predictors correlates strongly with agreement relative to AFDB across 1,595 proteins. The Pearson correlation is 0.971, and the Spearman correlation is 0.974 (Fig. S6).

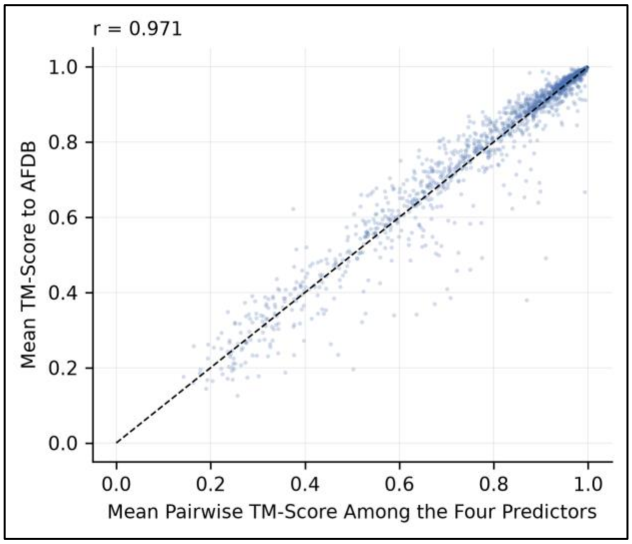


**Fig. S6.** Mean pairwise TM-score among the four HANSEN predictors is plotted on the horizontal axis against mean TM-score relative to the AFDB reference on the vertical axis. Each point represents one of 1,595 proteins. The dashed line is the identity line. Points remain close to this line across the full range. Agreement among the four predictors is therefore a reliable indicator of agreement with AFDB, even when the reference is not used.

#### 11. Agreement across the target-priority tiers

Agreement with AFDB is highest in the three prioritised tiers and lowest in the exploratory tier (Table S19). The tiers reported here are those assigned by the Platt-calibrated Target Priority Score that the resource now serves. For AlphaFold 3, the mean TM-score is 0.962 among the 50 high-priority targets, 0.956 among the 125 strong candidates and 0.928 among the 513 moderate candidates. The corresponding value among the 910 exploratory-tier proteins that have an AFDB counterpart is 0.766. The other three methods follow the same pattern. Mean model pLDDT follows the same order. For AlphaFold 3, it is 92.2 in the high-priority tier and 80.9 in the exploratory tier. Of the 140 proteins with a TM-score below 0.5 for all four methods, 139 belong to the exploratory tier. Table S20 reports the TM-score and lDDT for each method for the 50 highest-priority targets.

#### 12. Limitations

The AFDB models are predictions rather than experimental structures. Agreement with them therefore measures reproducibility rather than accuracy. The comparison with the seven experimentally determined *M. leprae* structures is reported in the main text. All comparisons in this note use only Cα atoms. Side-chain placement, rotamer geometry and hydrogen bonding are therefore outside the scope of the analysis. Secondary structure is assigned from Cα geometry, which identifies regular structure more freely than DSSP. None of the monomer models in this comparison contains a ligand, cofactor or metal, so these components are not considered. The differences between constructs in Table S10 arise because the AFDB release and the HANSEN models were built with different UniProt releases.

#### 13. Tables, data and code

Seventeen tables accompany this note in a workbook, with one worksheet for each table. They are numbered consecutively after the supplementary tables in the main text. Tables S11, S12, S17, S18 and S25 appear above. Tables S11 and S25 contain selected columns, and their complete versions are available in the workbook. The other twelve tables are available only in the workbook. Table S9 reports comparison coverage, while Table S10 lists the proteins whose modelled chain lengths differ between the two model sets. Table S13 reports the effect of limiting the analysis to the confident core. Table S14 reports agreement by chain length, Table S15 reports secondary structure and compactness, and Table S21 reports agreement by predicted localisation. Table S16 lists all discordant proteins, while Table S24 presents the named case studies. Table S19 reports agreement by target-priority tier, and Table S20 gives individual results for the 50 highest-priority targets. Table S22 contains per-protein results for the complete dataset, and Table S23 reports the validation of the TM-score implementation. Figs. S1 to S6 accompany the note.

Per-protein results for all 6,379 comparisons are provided in Table S22. We performed the comparison with the AFDB proteome archive UP000000806_272631_MYCLE_v6.tar and the HANSEN monomer set recorded in protein_characteristics.db.

### Supplementary Note SN2: Cross-species essentiality transfer, model selection and calibration

This note expands Section 2.7.1 of the main text. It sets out the limitations of transferring an essentiality classifier from *M. tuberculosis* to *M. leprae*, the comparison between the three feature sets that were considered, and the calibration of the reported probabilities.

#### SN2.1 Limitations of the cross-species transfer

Three limitations of the cross-species transfer should be borne in mind. First, ProteomeLM produces whole-proteome contextual embeddings, so proteins in a held-out fold contribute to the embeddings of proteins in the training folds; grouping folds by MMseqs2 clusters controls sequence similarity but not this contextual leakage, and the cross-validated AUROC of 0.84 is therefore an upper bound. Second, embedding proceeds in overlapping, genome-ordered windows, so the representation of a protein depends on its genomic neighbours; essential genes are frequently clustered in operons in *M. tuberculosis*, whereas gene order has been extensively rearranged in *M. leprae*, and the contextual signal is therefore unlikely to transfer intact between the two species. Third, the predicted essential fraction runs against expectation: the mean calibrated probability across the proteome is 0.089 and only 90 of 1,603 proteins are classified as essential, whereas a genome reduced to about 1,600 functional genes should if anything be enriched for essentiality relative to *M. tuberculosis*. This is more consistent with miscalibration of the base rate under cross-species transfer than with a biological finding, and the class counts should be read accordingly.

#### SN2.2 Model selection

Three feature sets were compared: ProteomeLM-L contextual features, ESM-C features alone, and their concatenation. Under homology-grouped cross-validation in *M. tuberculosis* the three are within one fold standard deviation of one another (AUROC 0.84 ± 0.01, 0.81 ± 0.01 and 0.83 ± 0.01), so cross-validation does not separate them. On transfer to the orthologue-aligned *M. leprae* set the ordering reverses sharply (0.78, 0.92 and 0.93). That set is not homology-separated: its 241 proteins share a median of 78% sequence identity with training proteins and are labelled by the essentiality of those proteins, whereas the cross-validation folds are grouped at 40% identity. Because those labels are the orthologue's own experimentally determined essentiality, assigning each protein the classification of its *M. tuberculosis* orthologue scores an AUROC of 1.000 by construction. The transfer column therefore rewards recovery of sequence similarity, which sequence-only features supply directly, and it cannot be used to rank the three feature sets. An earlier stratification of the transfer AUROC by orthologue identity carried only 7, 11 and 21 positives per band and is not informative in either direction. No cross-species ranking of the feature sets is therefore claimed.

#### SN2.3 Calibration of the reported probabilities

ProteomeLM-L-*leprae* was retained as the deployed model because the essentiality probability enters the Target Priority Score as a magnitude rather than a rank, which makes calibration the governing consideration, and because the probabilities are now calibrated. Against the *M. tuberculosis* essential-class base rate of 0.160 (625 of 3,902 labelled genes, at which the no-skill Brier score is 0.134), the raw probabilities scored a Brier score of 0.132, a Brier skill score of only +0.02. Platt scaling fitted on the held-out cross-validation probabilities reduces the Brier score to 0.100, a Brier skill score of +0.26, without altering the ranking (Fig. 3b of the main text). Discrimination is unchanged. An AUPRC of 0.54 against a base rate of 0.160 represents a 3.4-fold enrichment. All probabilities reported here, in the resource and in the Target Priority Score are the calibrated values. Calibration lowers the mean predicted essentiality across the proteome from 0.111 to 0.089, against an *M. tuberculosis* base rate of 0.160, and only five proteins now exceed a calibrated probability of 0.70. A genome reduced to about 1,600 functional genes might be expected to be enriched for essentiality relative to its parent, not depleted, and the model does not reproduce that expectation. The probabilities should therefore be read as a calibrated ordering for prioritisation and not as absolute estimates of indispensability in *M. leprae*.

### Supplementary Note SN3: Additional methodological notes

This note collects three methodological points supporting Sections 2.1, 2.9 and 2.11 of the main text.

#### SN3.1 Reconciliation of the proteome counts

The *M. leprae* strain TN genome encodes approximately 1,605 protein-coding genes in the current annotation release. The corresponding UniProt proteome contains 1,604 entries, one of which is the tmRNA-encoded proteolysis-tag peptide (UniProt V6B544), is not a protein-coding gene and was excluded, leaving 1,603 protein entries. These 1,603 UniProt entries resolve to 1,600 distinct loci, because three loci (ML1911, ML2219 and ML2569) are each represented by two UniProt entries. The 1,603 modelled and ranked proteins are a different set of the same size, and the agreement between the two totals is arithmetic rather than an identity. The ranked set comprises 1,599 loci that carry a UniProt accession together with four loci (ML1180, ML1181, ML1183 and ML2692) that were modelled from genomic translations and have no UniProt entry, and it excludes ML2428A (rpsW, UniProt P0A5D0), which the UniProt-keyed table registers as a locus in its own right and which carries a model but is not ranked. The figure of 1,599 given in Section 2.1 of the main text is therefore the number of ranked proteins carrying a UniProt accession and not a count of distinct gene names; both figures are correct once the two sets are distinguished.

**SN3.2 Derivation of the model-confidence values in Table S28**

Every value in Table S28 was independently re-derived from the B-factor field of all 7,122 model mmCIF files and reproduces the tabulated figures exactly, including the two rows in which the mean pLDDT and the percentage of models above 70 coincide to one decimal place; those coincidences are genuine and not a transcription error. As stated in Methods 4.4 of the main text, pLDDT is averaged over Cα atoms rather than over all atoms of each residue, and the two differ by about 2.6 units for AlphaFold 3 and Chai-1, whose mmCIF files carry per-atom rather than per-residue values. Separately, the all-model Boltz-2 row aggregates 564 co-folded assemblies with 140 constructed by rigid-body superposition of Boltz-2 monomers onto template assemblies; the latter inherit their confidence from the monomers and are not assembly predictions, and are identified as such in Table S2.

**SN3.3 Component ablation of the Target Priority Score**

A component ablation was run against the same validated set and returns two results that qualify the score. First, no single component matches the composite, but the annotation term comes close: alone it recovers the validated targets with an AUROC of 0.866 against 0.878 for the composite, while essentiality alone gives 0.737, model quality 0.656 and tractability 0.649. The composite is therefore justified relative to essentiality alone, which was the specific concern, but it is close to a re-expression of the annotation term. Second, and more seriously, the binding-site component does not discriminate at all. It takes the value 1.0 for every one of the 1,603 proteins, because the consensus pocket score saturates proteome-wide, so a term weighted 0.25 directly and a further 0.045 through tractability, together 29.5% of the Target Priority Score, is a constant offset that shifts every score equally and changes no ranking. Substituting the AF2BIND maximum binding probability, which does vary, lowers recovery (AUROC 0.804), so the constant is not concealing a better signal; but the term as published contributes nothing, and the score should be read as driven by annotation, essentiality and model quality. Rescaling the binding-site term so that it discriminates is the single most useful improvement available to this score and is noted as future work rather than claimed here.

**Supplementary figures and tables**

The main text is limited to ten figures and tables combined, so the items below are presented here. Figure and table numbering continues from Supplementary Note SN1 (Figs. S1 to S6) and Supplementary Workbook 2 (Tables S9 to S25).

(a) (b)

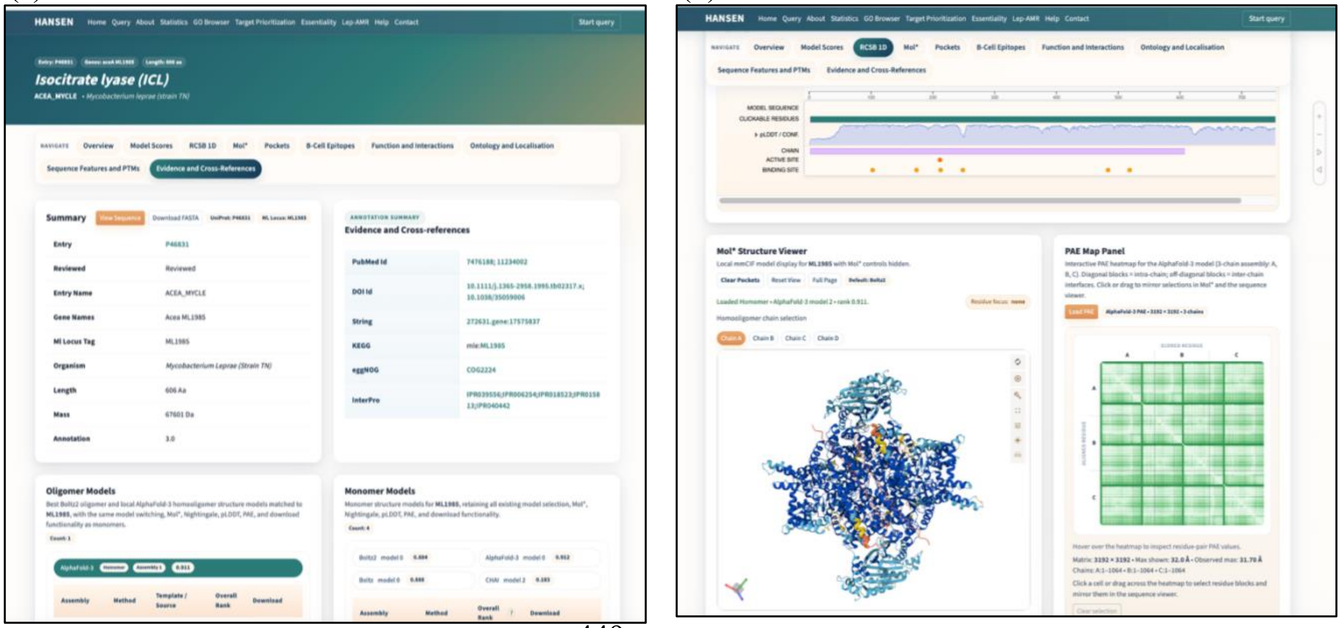


**Fig. S7.** Representative HANSEN protein page for isocitrate lyase (aceA, ML1985). (a) The page header, the summary of identifiers and annotation, the cross-references, and the oligomeric and monomeric model records with their ranking scores. (b) The residue-level sequence and feature viewer, the interactive Mol* viewer showing the predicted structure coloured by pLDDT, and the predicted aligned error map for the selected model.

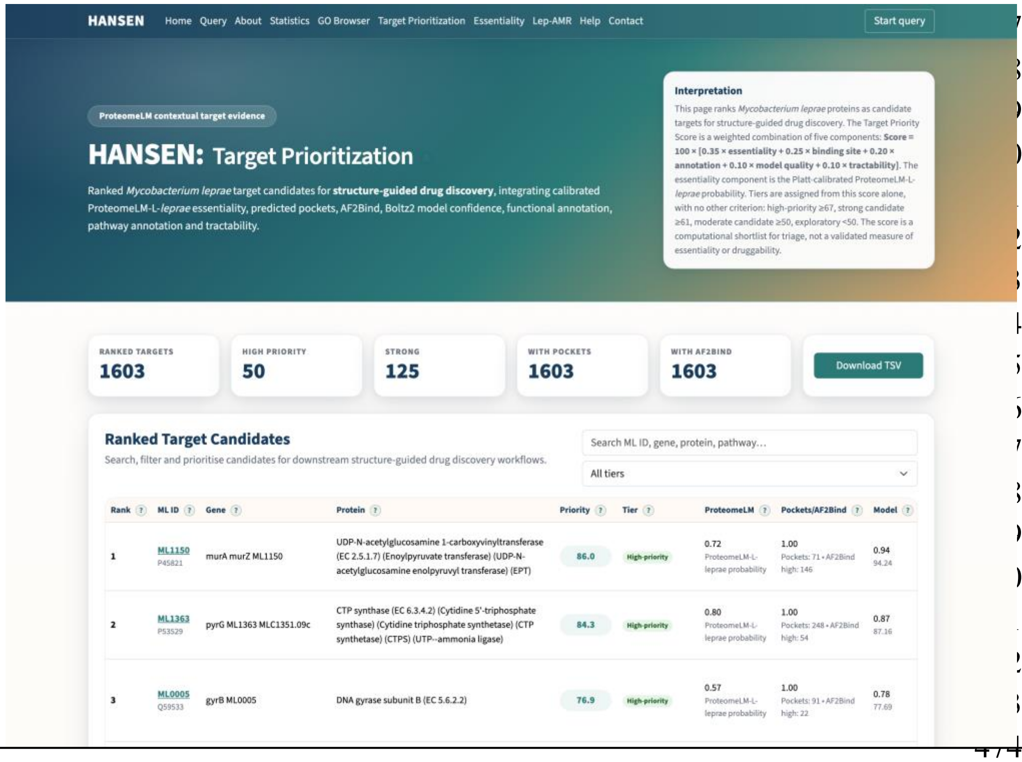


**Fig. S8.** HANSEN target-prioritisation browser. Proteins are ranked by the Target Priority Score and filtered by tier. Tier membership is determined by the score alone, with cut-points of 67, 61 and 50. The page shown here serves the Platt-calibrated Target Priority Score, under which 50 proteins are high-priority and 125 are strong candidates.

| **Scope** | **Method** | **Models (n)** | **Mean pLDDT** | **Median pLDDT** | **Mean pLDDT >70 (%)** | **Mean pLDDT >90 (%)** | **Residues pLDDT >70 (%)** |
|---|---|---|---|---|---|---|---|
| Monomeric | AlphaFold 3 | 1,603 | 85.3 | 89.2 | 88.9 | 46.0 | 87.2 |
| Monomeric | Boltz-1 | 1,599 | 81.2 | 86.2 | 81.2 | 32.6 | 82.4 |
| Monomeric | Boltz-2 | 1,599 | 80.6 | 85.7 | 80.6 | 29.3 | 81.7 |
| Monomeric | Chai-1 | 1,595 | 85.6 | 89.0 | 90.5 | 44.6 | 88.9 |
| All models | AlphaFold 3 | 1,625 | 85.3 | 89.1 | 88.9 | 45.7 | 86.9 |
| All models | Boltz-1 | 1,599 | 81.2 | 86.2 | 81.2 | 32.6 | 82.4 |
| All models | Boltz-2 | 2,303 | 82.0 | 86.0 | 84.6 | 31.8 | 81.2 |
| All models | Chai-1 | 1,595 | 85.6 | 89.0 | 90.5 | 44.6 | 88.9 |

**Table S28. Proteome-wide model confidence by modelling method,** reported separately for the monomeric models and for all structure models. Coverage columns give the percentage of models, or of residues, exceeding the indicated pLDDT thresholds. Monomeric scope is the like-for-like comparison because all four methods contribute a comparable number of monomeric models. Mean pLDDT is averaged over the Cα atoms of each model.

| Reference (ML id, gene, PDB) | Aligned residues | TM AlphaFold 3 | TM Boltz-1 | TM Boltz-2 | TM Chai-1 |
|---|---|---|---|---|---|
| ML0210 (ppa, 4ECP) | 159 | 0.98 | 0.99 | 0.98 | 0.98 |
| ML0380 (groES, 1LEP) | 81 | 0.82 | 0.88 | 0.87 | 0.80 |
| ML0482 (ruvA, 7OA5) | 180 | 0.77 | 0.21 | 0.34 | 0.73 |
| ML1382 (rpsA, 5IE8) | 89 | 0.88 | 0.87 | 0.87 | 0.88 |
| ML1806 (inhA, 2NTV) | 268 | 0.97 | 0.92 | 0.99 | 1.00 |
| ML2640 (2UYQ) | 274 | 0.96 | 0.97 | 0.98 | 0.97 |
| ML2684 (ssb, 3AFQ) | 105 | 0.91 | 0.90 | 0.90 | 0.89 |
| Mean | | 0.90 | 0.82 | 0.85 | 0.89 |

**Table S29. Accuracy of predicted models against the seven *M. leprae* experimental reference structures,** with TM-score reported for each method. Aligned residues are the sequence-matched Cα positions used for superposition against a single chain of the deposited structure. All seven structures were deposited before the training cut-offs of the four methods, so this comparison is a sanity check rather than an assessment of accuracy on unseen targets.

| Method pair | Proteins (n) | Mean TM-score | Median Cα-RMSD (Å) | pLDDT correlation (r) |
|---|---|---|---|---|
| AlphaFold 3 – Boltz-1 | 1,599 | 0.80 | 5.32 | 0.84 |
| AlphaFold 3 – Boltz-2 | 1,599 | 0.80 | 5.71 | 0.84 |
| AlphaFold 3 – Chai-1 | 1,595 | 0.81 | 4.64 | 0.84 |
| Boltz-1 – Boltz-2 | 1,599 | 0.83 | 3.83 | 0.96 |
| Boltz-1 – Chai-1 | 1,595 | 0.80 | 4.75 | 0.85 |
| Boltz-2 – Chai-1 | 1,595 | 0.80 | 5.14 | 0.85 |

**Table S30. Cross-method concordance among proteins modelled by more than one method.** TM-score and Cα-RMSD were calculated from sequence-matched, superposed Cα atoms. The correlation column shows the mean per-residue pLDDT correlation between each pair of methods. Values were derived from 9,582 pairwise comparisons.
